# TRPM5^+^ microvillous tuft cells regulate neuroepithelial intrinsic olfactory stem cell proliferation

**DOI:** 10.1101/2022.09.26.509561

**Authors:** Saltanat Ualiyeva, Evan Lemire, Caitlin Wong, Amelia Boyd, Evelyn C. Avilés, Dante G. Minichetti, Alexander Perniss, Alice Maxfield, Rachel Roditi, Ichiro Matsumoto, Nora A. Barrett, Kathleen M. Buchheit, Tanya M. Laidlaw, Joshua A. Boyce, Lora G. Bankova, Adam L Haber

**Author notes:** These authors contributed equally to this work (Saltanat Ualiyeva, Evan Lemire, Caitlin Wong).

## Abstract

The olfactory neuroepithelium serves as a sensory organ for odors and is part of the nasal mucosal barrier. Olfactory sensory neurons are surrounded and supported by epithelial cells. A subset of these, microvillous cells (MVCs), are strategically positioned at the apical surface but their specific functions are still enigmatic and their relationship to the rest of the solitary chemosensory cell family is unclear. Here, we establish that the larger family of MVCs comprises tuft cells and ionocytes in both mice and humans. Olfactory TRPM5^+^ tuft-MVCs share a core transcriptional profile with the chemosensory tuft family, prominently including the machinery for lipid mediator generation. Integrating analysis of the respiratory and olfactory epithelium, we define the unique receptor expression of TRPM5^+^ tuft-MVC compared to the Gɑ-gustducin*^+^* respiratory tuft cells and characterize a new population of glandular DCLK1*^+^* tuft cells. To establish how allergen sensing by tuft-MVCs might direct olfactory mucosal responses, we employed an integrated single-cell transcriptional and protein analysis. We defined a remodeling olfactory epithelial switch pathway with induction of *Chil4* and a distinct pathway of proliferation of the quiescent olfactory horizontal basal stem cell (HBC), both triggered in the absence of significant olfactory apoptosis. While the *Chil4* pathway was dependent on STAT6 signaling and innate lymphocytes, neither were required for HBC proliferation. HBC proliferation was dependent on tuft-MVCs, establishing these specialized epithelial cells as both sensors for allergens and regulators of olfactory stem cell responses. Together our data provide high resolution characterization of the nasal tuft cell heterogeneity and uncover a novel mechanism by which TRPM5^+^ tuft cells direct the olfactory mucosal response to allergens.

**One Sentence Summary:** We identify the enigmatic TRPM5^+^ olfactory microvillous cells as tuft cells, and show their functional role as regulators of olfactory stem cell proliferation in response to environmental signals.

## INTRODUCTION

The nasal cavity is the first site of contact of inhaled air within the respiratory tract. It is divided by the septum into two air passages and an intricate system of turbinates and sinus cavities where inhaled air is humidified, its temperature equilibrated to that of the lower airways and large particulate matter is trapped. The lining of the nasal cavity contains two functionally distinct but anatomically overlapping mucosal compartments: respiratory epithelium and olfactory neuroepithelium, overlying a dense network of submucosal glands. The olfactory neuroepithelium is a unique neuronal organ comprised of olfactory sensory neurons (OSNs) surrounded by specialized epithelial cells. The OSNs continuously regenerate from a population of proliferating progenitor cells called globose basal cells (GBCs). A separate subset of stem cells, horizontal basal cells (HBCs), are quiescent and serve as a reserve population, and can be activated by profound damage with loss of neuroepithelial structure (1, 2). Interspersed in both the respiratory and olfactory mucosa are immune cells. Thus, the nose represents an extraordinary site of close interaction of epithelial and progenitor cells, secretory glands, the immune and nervous systems (3). Recent mechanistic studies have highlighted the critical contribution of the nasal mucosal response to the development of systemic immunity (4, 5), and the distinct mucosal immune responses in the olfactory and respiratory nasal epithelium (6).

Respiratory epithelial cells play a central role in the recognition of danger signals and their mediators direct the polarity of the airway immune response (7). While epithelial cell damage from viruses, bacteria and toxins initiates a protective inflammatory response, receptor-driven allergen activation of epithelial cells skews the immune response to a type 2 “remodeling” effect. Similar to their respiratory counterparts, olfactory epithelial cells play a role as immune modulators. The olfactory sustentacular cells recently took center stage in mucosal immunology as the major expression site of the dominant SARS-CoV-2 entry receptor ACE2. Upon SARS-CoV2 infection sustentacular cells rapidly turn on proinflammatory signals including production of interleukins (IL-1β) and monocyte-recruiting chemokines (CCL2-5)(8). In addition to sustentacular cells, two subsets of microvillus cells (MVCs) present unique epithelial cells with emerging functions in mucosal immune responses (9).

The olfactory MVCs consist of two distinct subsets (10). The smaller pear shaped MVCs lining the apical surface of the olfactory neuroepithelium are distinguished by the expression of the Transient Receptor Potential Cation Channel Subfamily M Member 5 (TRPM5), a taste receptor-linked calcium-activated monovalent cation channel (10, 11). A distinct subset of MVCs with broad cell body and a slender cytoplasmic process extending to the basement membrane is distinguished by the expression of the type 3 IP3 receptor (IP3R3) and the 5’ exonucleotidase CD73 while they lack expression of TRPM5 (10, 12-15). Both TRPM5^+^ and TRPM5^−^ MVCs are derived from a common kit^+^ progenitor but differ in their differentiation trajectory (14, 16). The two subsets of MVCs are also shown to have distinct functions. IP3R3^+^ MVCs are a source of the neuropeptide Y and direct the activation and proliferation of GBCs (15, 17). TRPM5^+^ MVC are linked to impaired olfactory-guided behaviors after extended exposure to strong odorants (18). TRPM5^+^ MVCs activation with strong odorants, bacterial lysate and cold temperatures leads to acetylcholine release and downstream calcium flux in olfactory sustentacular cells suggesting that TRPM5^+^ MVCs sense environmental signals and their activation signals direct the integrated response in the olfactory epithelium (19, 20).

Interestingly, TRPM5^+^ MVCs also express the acetylcholine-producing enzyme choline acetyltransferase (ChAT) (19). TRPM5 and ChAT notably also mark tracheal tuft (also known as brush) cells (21, 22), intestinal tuft cells (23, 24), and the related nasal solitary chemosensory cells (SCC)(25, 26), which suggests shared activation pathways between tuft cells and TRPM5^+^ MVCs. TRPM5^+^ MVC development depends on Pou2f3 (27), a transcription factor required for tuft cell development in all mucosal compartments (28–30). Tuft cells are the dominant epithelial source of IL-25 and a relevant source of innate cysteinyl leukotrienes (CysLTs)(31–36). Tuft cell activation by metabolites derived from microbiota and allergens directs the activation of type 2 innate lymphoid cells (ILC2) for IL-13 driven type 2 inflammation and stem cell activation (30, 31, 34, 35, 37). However, other defining features of tuft cells such as the expression of taste receptor type I and II families and the taste signaling G protein Gα gustducin (38, 39) are low in TRPM5^+^ MVCs (10, 11, 27, 40). Our previous studies demonstrated that ChAT-eGFP^+^ nasal epithelial cells with morphology and transcriptional features of TRPM5^+^ MVCs are directly activated by allergens and generate cysteinyl leukotrienes (32, 41). Whether the TRPM5^+^ MVCs engage the same tuft-ILC2 circuit of inflammation and stem cell proliferation in the olfactory mucosa after allergen inhalation is not known. As the TRPM5^+^ MVCs are notably more abundant at homeostasis than the rare tuft cells, their functions might provide insight into the role of hyperplastic tuft cells such as those found in the recovery phase after viral infections and bleomycin injury of the lung (42, 43).

Here, we used ChAT-eGFP mice to enrich TRPM5^+^ MVCs from the olfactory and SCCs from the respiratory epithelium for single-cell RNA-sequencing (scRNAseq) analysis demonstrating that TRPM5^+^ MVC, SCCs and a third “intermediate” population of ChAT-eGFP^+^ nasal epithelial cells all belong to the tuft cell family. Although each population can be defined by unique markers, they share a core transcriptional profile including *Trpm5, Pou2f3, Chat, Avil* and *Ltc4s*. The transcriptionally distinct population of TRPM5^−^ MVCs shares multiple features with pulmonary ionocytes (*Cftr, Foxi1, Ascl3, Coch*). We applied supervised machine learning approaches to accurately assess the transcriptome-wide similarity of detected cell-types to known subtypes in the trachea, which confirmed the identity of TRPM5^+^ MVCs as tuft cells, and of TRPM5^−^ MVCs as ionocytes. We assembled the largest single-cell atlas of the mouse nasal mucosa (containing 63,125 cells) including analysis of the response to the inhaled mold allergen *Alternaria*. We found that allergen inhalation caused a marked proliferation of HBCs in the olfactory mucosa, even in the absence of profound damage to the OSNs or accessory epithelial cells. Strikingly, this allergen-induced stem cell proliferation was abolished by genetic deletion of tuft cells using *Pou2f3^−/−^*mice, demonstrating that tuft cells are critical for olfactory stem cell activation. We further used genetic ablation of lymphocytes (*Il7r^−/−^*) and IL-13 signaling (*Stat6*^−/−^) to show allergen-induced stem cell proliferation does not depend on type 2 inflammatory pathways. Taken together, our data define extensive heterogeneity of sensory epithelial cell-types in the nasal mucosa and define the functional consequences of their effector programs for stem cell proliferation.

## RESULTS

### Mouse olfactory TRPM5^+^ MVCs are tuft cells and IP3R3^+^ MVCs are ionocytes

First, to define the distribution of mouse olfactory and nasal respiratory chemosensory ChAT-eGFP^+^ cells by flow cytometry (FACS), we separated olfactory from respiratory mucosa along the anterior portion of the olfactory turbinates and obtained single-cell suspensions from each compartment (**Fig. S1A**). The olfactory mucosa contained the majority of ChAT-eGFP^+^ cells (19) (69%) compared to 31% derived from the respiratory mucosa (**Fig. S1B-F**). ChAT-eGFP^+^ cells were abundant in the olfactory mucosa (22% of the epithelial cell adhesion molecule (EpCAM)^high^ cells and 2% of live cells) but were also surprisingly prevalent in the respiratory mucosa (10% of EpCAM^high^ cells and 1% of live cells) (**Fig. S1B-E**). Our previous analysis of the flow cytometric features of ChAT-eGFP^+^ nasal epithelial cells defined two subsets: globular (FSC/SSC^low^) and elongated (FSC/SSC^high^) (32). Bulk RNAseq analysis of sorted cells of each subset suggested they correspond to the historical description of olfactory MVCs (globular) and respiratory SCCs (elongated) (32). Surprisingly, here we found that the olfactory and respiratory mucosa had similar distribution of globular vs elongated ChAT-eGFP^+^ cells suggesting that the abundant MVCs, previously described exclusively in the olfactory mucosa, are also be found in the respiratory epithelium albeit in lower numbers (**Fig. S1G-H**). Elongated ChAT-eGFP^+^ cells were also found in both respiratory and olfactory mucosa. They were extremely rare in both locations accounting for less than 500 cells per mouse (**Fig. S1H**). To validate the separation of olfactory and respiratory mucosa we used aquaporin 4 as a distinguishing marker between olfactory and vomeronasal sensory neurons (VSNs) in rodents (44) (**Fig. S2A-C)**. Although the VNO is an olfactory organ, it is located in the anterior part of the nose surrounded by respiratory epithelium (**Fig. S1A**). We found nearly all *Aqp4^+^* neurons in our single cell preparations were derived from the respiratory portion (**Fig. S2A, C**), where the aquaporin 4 positive VSNs are located (**Fig. S2B**), validating our separation of tissue origin.

To characterize the subsets of nasal ChAT-eGFP^+^ cells transcriptionally, we compared them to EpCAM^high^ nasal epithelial cells from each of olfactory and respiratory epithelia by scRNAseq (**Fig. 1A** and **S1A**). To ensure sufficient SCC cell numbers, we enriched for the scarce elongated ChAT-eGFP^+^ cells so they represented one third of the cells submitted for scRNAseq, while another third was the ChAT-eGFP^+^ globular cells and the remainder were EpCAM^high^ GFP^−^ cells (**Fig. S1I)**. Dimensionality reduction and unsupervised clustering (**Methods**) identified a subset of cells defined by known tuft cell markers (**Table S1** and **S2**) that clustered distinctly from all other EpCAM^high^ cells from the respiratory and olfactory mucosa (**Fig. 1A**). Transcriptome-wide machine learning analysis confirmed that these are indeed highly similar to tracheal tuft cells, with 94% of the cells in the TRPM5^+^ MVC cluster being classified as tuft cells by a random forest classifier (**Methods**) trained on previously published single-cell data from the tracheal epithelium (36) (**Fig. 1B**). Besides tuft cells, we identified a second population of differentiated epithelial cells negative for *Chat* and *Trpm5* but positive for *Itpr3* and *Ascl3*, consistent with descriptions of TRPM5^−^ MVCs (15, 45, 46). These cells expressed high levels of *Cftr, Coch* and *Foxi1*, and the random forest mapped 96% to tracheal ionocytes (**Fig. 1B**), unambiguously identifying them as nasal ionocytes (**Fig. 1B**).

**Fig. 1.**
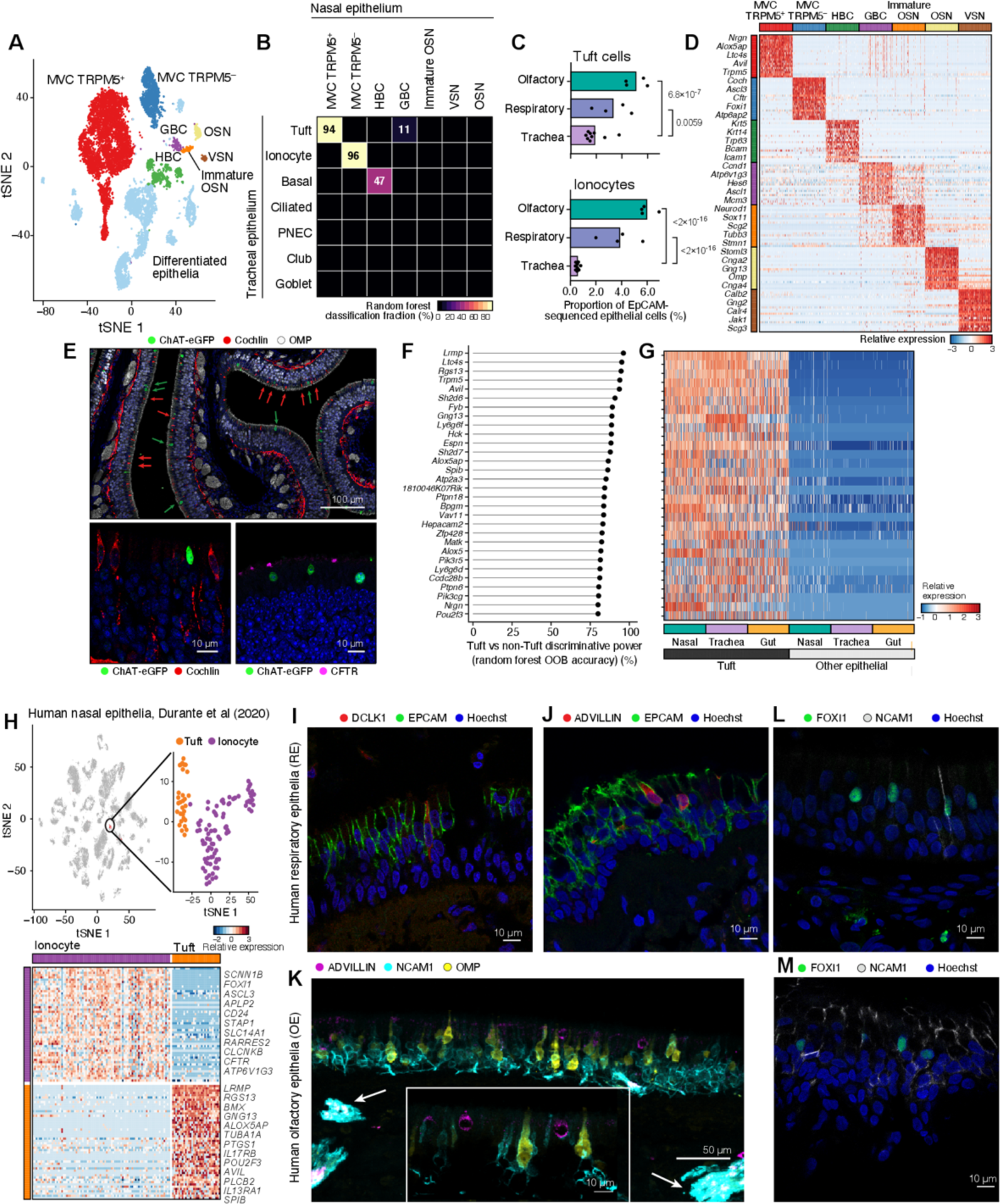
Tuft cells and ionocytes comprise two MVC types of the olfactory epithelium in mice and humans. (**A-G**) Single cell sequencing (**A-D** and **F-G**) and histology (**E**) of mouse nasal MVCs. (**A**) tSNE embedding of 13,052 nasal epithelial cells (*n*=4 mice), colored and labeled by unsupervised clustering. (**B**) Heatmap shows the fraction of cells (color bar) of each nasal epithelial cell type (columns, types as in **A**) classified as each tracheal epithelial cell type (rows) using a random forest. (**C**) Fraction (x-axis) of ionocytes in all epithelial cells from each mouse (points), compared to tracheal epithelium. (**D)** Heatmap shows relative expression (row-wise z-score of log_2_(TPM+1)), (color bar) of top 50 genes (rows) specific to each cell type (FDR<0.05) in our dataset; two selected genes are shown. (**E**) Cross-section of the nasal olfactory epithelium stained for ChAT-eGFP, cochlin, OMP (top image), ChAT-eGFP and cochlin (lower left image), and ChAT-eGFP and CFTR (lower right image). Hoechst is used to mark nuclei in all images. (**F**) Top 30 pan-tissue tuft cell marker genes (y-axis) ranked by out of bag (OOB) accuracy (x-axis) from random forest classification. (**G**) Heatmap shows relative expression of top tuft cell marker genes (ordered as in **F**) in tuft cells and other epithelial cells across three tissues: nasal, trachea, and gut. (**H-M**) Single cell sequencing (**H**) and histology (**I-M**) of human nasal MVCs. (**H**) tSNE embeddings (top) of 3,528 olfactory epithelial, immune and neuronal cells (54) showing new re-analysis by unsupervised clustering revealing a cluster of MVCs containing tuft cells and ionocytes (inset), and heatmap (bottom) of differentially expressed genes (FDR<0.05, rows) between the 116 identified MVCs (54). Selected marker genes are labeled. (**I-M**) Immunofluorescence of respiratory epithelium (**I, J, L**) or olfactory neuroepithelium (**K, M**) from ethmoid sinus or superior turbinates of patients with CRS (*n*=3) or controls without sinus disease (**K-M**) stained for DCLK1 (**I**) or advillin (**J, K**) in combination with EpCAM (**I-J**) to mark epithelial cells or with NCAM1 (**K-M**) to mark OSNs. Arrows in **K** indicate olfactory axon bundles in the submucosa.

Although both tuft cells and ionocytes still represent a minority of epithelial cells, they appeared markedly more abundant in the nose than their reported concentration in the trachea (36). To characterize their actual distribution in the nose, we evaluated the numbers of tuft cells and ionocytes in a non-enriched scRNAseq dataset from the nose (to be introduced in **Fig. 3**) and compared to published tracheal scRNAseq data (36). Examining this non-enriched dataset, we found that tuft cells comprise 5.1% (95% confidence interval (CI): 4.9-5.7) of EpCAM^+^ cells from the olfactory epithelium, 2.7-fold more common than in the trachea (**Fig. 1C**). Ionocytes comprise 5.6% (95% CI: 5.3-6.5) of EpCAM^+^ cells from the olfactory epithelium, 18.7-fold more common than in the trachea (**Fig. 1C**). Nasal tuft cells were defined by the expression of the canonical markers defined by our previous scRNAseq analyses of the trachea and intestine: the taste transduction channel *Trpm5,* Ca^2+^-regulated protein *Avil,* and eicosanoid biosynthetic enzymes *Alox5ap* and *Ltc4s* (**Fig. 1D** and **Table S2**). Immunofluorescence of cross sections of nasal mucosa confirmed that both cochlin^+^ and CFTR^+^ ionocytes and ChAT-eGFP^+^ tuft cells are abundant in the olfactory mucosa (**Fig. 1E**) and ionocytes are found in lower numbers in the respiratory epithelium (**Fig. S2D**). The remainder of the EpCAM^high^ cells were comprised of differentiated epithelial cells, basal cells, and olfactory major protein (OMP) expressing sensory neurons (**Fig. 1A**, **D**).

TRPM5^+^/ChAT^+^ cells in the olfactory and respiratory mucosa of the nose are historically defined as distinct epithelial subsets – olfactory MVCs and respiratory SCCs, largely based on the absence of taste receptors and transduction machinery in MVCs (11, 47). However, we find that *Trpm5^+^*/ChAT^+^ cells derived from both the olfactory and respiratory mucosa share a core transcriptional profile defined by transcripts involved in taste transduction (*Trpm5, Gng13*), multiple calcium signaling molecules (*Lrmp, Pik3cg, Hck, Vav1, Matk, Pik3r5*), the transcription factors (TFs) *Spib* and *Pou2f3* and the cysteinyl leukotriene (CysLT) biosynthetic enzymes *Alox5, Alox5ap* and *Ltc4s* (**Fig. 1F**). This core effector profile is shared by tuft cells in the trachea (36) and intestine (35) defined by our previous single-cell analysis (**Fig. 1G**). Thus, tuft cells in all mucosal compartments are distinguished by the expression of the necessary machinery to rapidly generate Ca^2+^-signaling dependent mediators.

### Human olfactory TRPM5^+^ MVCs are tuft cells and IP3R3^+^ MVCs are ionocytes

Having characterized the subsets of respiratory and olfactory mouse nasal tuft cells, we sought to determine whether their profile and distribution is mirrored in the human nose. We reanalyzed existing scRNAseq datasets and immunophenotyped nasal samples from the respiratory and olfactory mucosa. Human SCCs in the respiratory epithelium share many of the reported features of mouse SCCs including taste receptors and IL-25 expression, and their numbers are higher in patients with fungal and chronic rhinosinusitis compared to healthy controls (48–50). To define the human SCC transcriptional profile, we reanalyzed scRNAseq profiles of nasal epithelial cells derived from sinus surgeries of patients with chronic rhinosinusitis with or without polyposis (**Fig. S2E, Table S4, Methods**) (51) and confirmed that 13 of the 19,196 cells profiled were indeed nasal tuft cells. They were identifiable by their specific expression of pan-tuft cell markers *LRMP, RGS13, AVIL* (**Fig. S2F**), taste transduction protein *PLCB2*, and transcripts for eicosanoid enzymes *ALOX5* and *PTGS1*. By histology, we found rare DCLK1^+^ spindle shaped cells in the respiratory epithelium of ethmoid sinus and superior turbinates derived from patients (*n*=3) with chronic rhinosinusitis using both DCLK1 and advillin immunostaining (**Fig. 1I-J**).

Human MVCs have been described by anatomical studies of olfactory mucosa, but it is not known whether these elongated cells represent tuft cells, ionocytes or both (52, 53). To determine whether they share the features of mouse MVCs, we reanalyzed a recent scRNA-seq dataset of human olfactory mucosa by Durante *et al* (54). Focused analysis of the cluster of 116 cells annotated as MVCs identified two subsets (**Fig. 1H**), and differential expression analysis identified one subset of human MVCs with highly similar transcriptional makeup to mouse tuft cells, with distinct expression of eicosanoid pathway components *ALOX5AP* and *PTGS1,* the IL-25 receptor *IL17RB*, the pan-tuft cell markers *AVIL* and *LRMP* and the tuft cell transcription factor *POU2F3* (**Fig. 1H, Table S4**). To locate human tuft cells in olfactory biopsies, we used *NCAM1* as a specific marker (55) expressed at high mRNA levels in both mature and immature human OSNs **(Fig. S2G)**. NCAM1 antibody marks human olfactory neuroepithelium including the axons of mature OSNs in proximity to OMP^+^ OSNs (**Fig. 1K**). In the olfactory neuroepithelium, advillin^+^ cells with the globular morphology of olfactory TRPM5^+^ MVCs were interspersed between OSNs suggesting a shared morphology with the globular mouse TRPM5^+^ MVCs (**Fig. 1K**). Finally, we confirmed that human tuft-MVCs express *ALOX5, ALOX5AP,* and *LTC4S*, required for CysLT generation (**Fig. S2G**) providing a clear functional connection to mouse tuft cells.

In addition to tuft cells, our re-analysis of scRNAseq data also identified a cluster of 79 ionocytes in the human respiratory epithelium (**Fig. S2E-F**) and 86 ionocytes in the olfactory enriched human mucosa (**Fig. 1H**). The human nasal ionocytes shared the transcriptional profile of human pulmonary ionocytes and mouse pulmonary and nasal ionocytes with unique expression of the ion channels *CFTR, CLCNKB*, the transcription factors *FOXI1* and *ASCL3*, and the vacuolar ATP-ase *ATP6V1G31*. We validated the presence of ionocytes in human respiratory and olfactory nasal tissue using expression antibody staining against FOXI1 (**Fig. 1L-M**). In summary, we confirmed that the human tuft cells and ionocytes represent two distinct subsets of the previously described human MVCs in the olfactory mucosa. Human tuft cells are heterogeneous in the nose similar to mouse tuft-MVCs and tuft-SCCs but share a core transcriptome. Ionocytes, a more homogenous population of specialized epithelial cells, are found in the olfactory mucosa as previously reported (36), but also in high numbers in the respiratory mucosa.

### Three subsets of mouse nasal tuft cells correspond to MVCs, SCCs and a new population of glandular tuft cells

Our scRNAseq evaluation demonstrated that all *Chat^+^Trpm5^+^* nasal epithelial cells share the tuft cell transcriptional profile. To reconcile this data with the literature that defined olfactory MVCs as distinct from respiratory SCCs and other chemosensory cells, we next sought to dissect their heterogeneity at the single-cell level. Unsupervised clustering identified three groups of tuft cells (**Fig. 2A**), and we used differential expression tests to define the unique markers of each cluster (**Fig. 2B, Table S3**). We confirmed that all express the pan tuft cell signature defined by Nadjsombati *et al.* (33) (**Fig. S3A)**. Each subtype expressed distinct receptors (**Fig. 2B**), such as those for norepinephrine: beta-2-adrenergic receptor (*Adrb2*), alpha-2-adrenergic receptor (*Adra2a*), beta-1 adrenergic receptor (*Adrb1)* in the tuft-MVC, tuft-gland, and tuft-SCC subtypes (rationale for nomenclature is below) respectively, raising the intriguing possibility of distinct interactions with norepinephrine secreting sympathetic neurons (56).

**Fig. 2.**
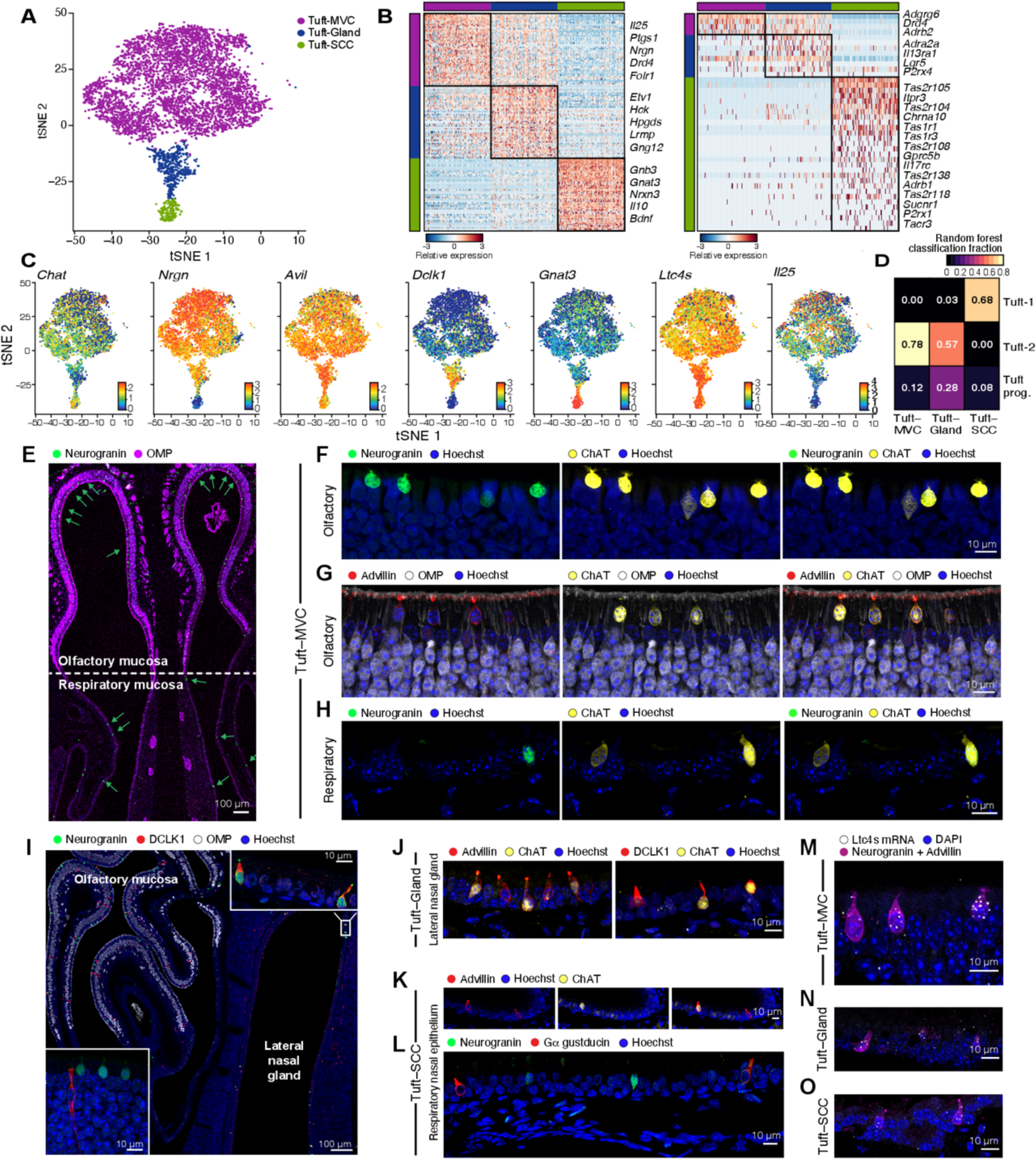
Single-cell analysis identifies nasal tuft cell heterogeneity in mice. (**A**) tSNE embedding of 5,594 tuft cells derived from ChAT-eGFP mice colored by unsupervised clustering into subsets. (**B**) Heatmaps show the expression level (row-wise z-score of log_2_(TPM+1), color bar) of marker (left) and receptor genes (right) specifically expressed (FDR<0.01) by each subset (color bars). Adrenergic and cholinergic receptors are highlighted in bold. (**C**) tSNE plots display expression in log_2_(TPM) + 1 (color bar) for shared tuft cell markers and subtype markers. (**D**) Heatmap shows the fraction of cells (color bar) of each type (rows) of each cell type (columns) using a random forest classifier trained on previously published scRNA-seq profiling of the tracheal epithelium (36). (**E-L**) Immunolocalization of tuft cell subsets. (**E**) Cross section through the anterior nasal cavity at the level of the nasal septum captures the olfactory epithelium (top) and respiratory epithelium (bottom). OMP reactivity delineates the OSNs and OSN axons co-stained with neurogranin. Arrows point at neurogranin positive cells. (**F-H**) Tuft-MVC expression of ChAT-eGFP, neurogranin (**F, H**) and advillin (**G**) in the olfactory (**F-G**) and respiratory (**H**) mucosa. (**I**) Cross section through the posterior section of the nasal cavity captures the olfactory epithelium (top left), respiratory epithelium (bottom center) and lateral nasal gland (right). OMP reactivity delineates the OSNs co-stained with neurogranin and DCLK1. Inserts show high magnification of the olfactory mucosa (lower left) and the respiratory epithelium lining the lateral nasal gland (top right). (**J**) Expression of ChAT-eGFP and advillin (left) or DCLK1 (right) in tuft cells lining the epithelium of the lateral nasal gland. **K-L**. Tuft-SCC expression of ChAT-eGFP and advillin (**K**), neurogranin, Gα gustducin (**L**). Hoechst was used for nuclear staining. (**M-O**) *In situ* hybridization with RNAscope probe for *Ltc4s*. Tuft cells were distinguished based on immunoreactivity for advillin and neurogranin. Representative images of tuft-MVC (**M**), tuft-Gland (**N**) and tuft-SCC (**O**) Nuclear staining with DAPI.

The dominant population of tuft cells (**Fig. 2A**) was largely derived from the olfactory mucosa (**Fig. S3B-C**), and was defined by the high and ubiquitous expression of tuft cell markers *Pou2f3, Trpm5, Chat* (the original markers of MVCs (11, 19, 27, 38), as well as the pan-tuft marker *Avil,* the cyclooxygenase enzyme *Ptgs1* (**Fig. 2B-C** and **S3D-E**). In whole mount stains of the olfactory mucosa, we found that ChAT-eGFP^+^ cells were abundant and displayed a globular body and a single very short process ending with short brushes at the apical surface, identical to the morphology of TRPM5^+^ MVCs (47) (**Fig. S3F-G**). These cells expressed low levels of limited numbers of taste receptors and low levels of the transcript for the taste receptor associated protein Gɑ-gustducin *Gnat3* (**Fig. 2C**). Albeit low, *Gnat3* as well as the related *Gnb3* was detectable in ∼50% of tuft-MVCs (**Fig. S3E**) consistent with other reports (40). Notably, TRPM5^+^ MVCs express high levels of the key eicosanoid biosynthetic enzymes *Hpgds, Alox5, Alox5ap* and *Ltc4s* (**Fig. 2C** and **S3H**) as well as the tuft cell-specific cytokine *Il25* (**Fig. 2B-C** and **S3H**). Since these cells share the morphology of MVCs but the core transcriptional profile of tuft cells, we refer to these small, globular *Chat*^+^*Trpm5*^+^ cells as tuft-MVCs. We identified a new marker of the TRPM5^+^ tuft-MVCs: *Nrgn*, encoding neurogranin, a postsynaptic neuronal protein in the calpacitin family that preferentially binds to the Ca^2+^-free form of calmodulin (57) (**Fig. 2B-C**). We confirmed the expression of neurogranin at the protein level (**Fig. 2E-F**), and its absence in tuft cell-deficient *Pou2f3^−/−^* mice, validating neurogranin as a true tuft cell marker (**Fig. S4A-B**). Nearly all neurogranin^+^ cells were also ChAT-eGFP^+^ but negative for Gɑ gustducin protein in the olfactory mucosa (**Fig. 2F** and **S4C-D**). Scattered neurogranin^+^ cells were also detectable in the respiratory epithelium (**Fig. S4B, D-E**). The ChAT-eGFP^+^ MVCs were positive for the pan-tuft cell marker advillin at the protein level (**Fig. 2G**). Finally, TRPM5^+^ tuft-MVCs had very low levels of the transcript (*Dclk1*) and protein DCLK1, commonly used to identify tuft cells in the intestine and trachea (30, 31, 58) (**Fig. 2C**, **I)**. Notably, besides tuft-gland cells, DCLK1 also marked epithelial cells with ductal morphology in the olfactory mucosa (**Fig. 2I inset**) indicating that in the nose DCLK1 is *not* a tuft cell-specific marker. Comparison to the tracheal scRNAseq-defined tuft cell subsets (5) using a machine learning classifier showed highest similarity to the previously identified tracheal eicosanoid-enriched tuft-2 cells (36) (**Fig. 2D**).

A second distinct but rare population of tuft cells were derived exclusively from the respiratory epithelium (**Fig. 2A** and **S3B-C**). These cells were enriched for transcripts that define the elongated nasal respiratory SCCs, including type 1 (*Tas1r1, Tas1r3*) and type 2 (*Tas2r104, Tas2r105, Tas2r108, Tas2r118*, and *Tas2r138*) taste receptors, and ubiquitous and high expression of gustducin components *Gnat3* and *Gnb3* (25, 38, 48, 59-61) (**Fig. 2B-C** and **S3E**). As expected, these rare respiratory tuft cells have a spindle-shaped morphology (**Fig. S3F**) and are also enriched for the core tuft cell genes (**Fig. 2C, S3A, D-E, H**). Because of this morphology and marker expression characteristic of SCCs, combined with the transcriptional profile of tuft cells, we refer to them as tuft-SCCs. Although with distinct receptor expression, tuft-SCCs expressed mediator transcripts *Il25, Alox5, Alox5ap, Ltc4s, Ptgs1,* and *Hpgds* highly and ubiquitously (**Fig. 2C**, **S3H**). As we found for tuft-MVCs, the tuft-SCC expression of *Dclk1* was very low (**Fig. 2C**). The tuft-SCCs were most similar to the taste receptor-enriched tracheal tuft-1 with 68% classified as tuft-1 (**Fig. 2D**). Histologically, tuft-SCCs were restricted to the respiratory epithelium and were largely negative for neurogranin (**Fig. S4D-E**) but marked by ChAT-eGFP, advillin and Gɑ-gustducin (**Fig. 2K-L** and **Fig. S4E-F**). Surprisingly, tuft-SCCs accounted for the dominant expression of the neuropeptide bone-derived neurotrophic factor (*Bdnf*) and the cytokine *Il10* in the epithelium at homeostasis (**Fig. 2B**) suggesting a possible immunomodulatory role.

Finally, we identified an intermediate population of tuft cells that shared features with both tuft-MVC and tuft-SCCs yet was transcriptionally distinct (**Fig. 2A-B**). Among the shared features were mediator transcripts *Il25, Alox5, Alox5ap, Ltc4s, Ptgs1,* and *Hpgds* (**Fig. 2C**, **S3H**). This subset expressed the highest level of *Il13ra1* (**Fig. 2B**), which is responsible for tuft cell proliferation in the intestine in the setting of type 2 inflammation (34), and consistently, they were most similar to the tuft cell progenitor population in the trachea (**Fig. 2D**). They were enriched for *Ly6d* and *Ly6e,* transcripts associated with breast and prostate glandular cancers (62–64) (**Table S3**). These ChAT^+^/advillin^+^ tuft cells were also positive for neurogranin, Gɑ-gustducin and DCLK1, and thus correspond to the intermediate population (**Fig. 2C, I-J, S3E** and **S4G-H**). This distinct population of tuft cells expressing both tuft-MVC and tuft-SCC markers was dominantly found in the lateral nasal gland – a large serous glandular structure unique to the nasal cavity (**Fig. 2I**). Thus, based on their transcriptional profile and their anatomical location in the lateral nasal gland we refer to them as tuft-gland. To confirm the morphological and anatomical characteristics of the nasal tuft cell subsets, we sorted elongated (FSC^high^) and globular (FSC^low^) tuft cells from the olfactory and respiratory epithelium for bulk RNAseq. We applied a support-vector machine regression deconvolution approach CIBERSORTx (65) to infer the proportions of the newly defined tuft cell subsets. The estimated compositions suggested that tuft-MVCs represent the majority of globular (FSC^low^) nasal ChAT-eGFP^+^ cells, and also that tuft-gland cells were equally represented in elongated (FSC^high^) and globular (FSC^low^) gates, and likely of intermediate morphology (**Fig. S3I**).

To confirm the shared expression profiles in the different morphologic subsets of nasal tuft cells, we used *in situ* hybridization for *Ltc4s* mRNA. Consistently with our scRNA-seq data (**Fig. 2C**), *Ltc4s* mRNA was ubiquitously found in all subsets of nasal tuft cells: tuft-MVCs in the olfactory mucosa (**Fig. 2M**), tuft-gland in the lateral nasal gland (**Fig. 2N**) and tuft-SCCs in the respiratory epithelium (**Fig. 2O**). We also characterized the vomeronasal organ (VNO) ChAT-eGFP^+^ cells as positive for Gα-gustducin (59) but negative for neurogranin and DCLK1 making them most similar to tuft-SCC despite their location surrounded by OMP^+^ neurons (**Fig. S4I-K**).

Taken together, these data define and characterize three subsets of nasal tuft cells with shared and distinct transcriptional, morphological, and anatomical features. All three subsets of nasal tuft cells shared expression of *Trpm5,* the acetylcholine generating enzyme *Chat,* the tuft cell cytokine *Il25* and transcripts of the eicosanoid generating cascade *Ptgs1, Hpgds, Alox5, Alox5ap* and *Ltc4s.* The three subsets could be distinguished based on their expression of neurogranin (tuft-MVC and tuft-gland), protein expression of Gɑ-gustducin (tuft-SCC and tuft-gland) and DCLK1 (only tuft-gland) as well as the distinct receptor expression of taste and adrenergic receptors.

### Characterization of the mouse nasal mucosal immune system

As we and others have shown, tuft cells are epithelial sensors of allergens that regulate local mucosal responses (31, 32, 41, 66). We therefore sought to comprehensively profile the nasal mucosal immune and neuroepithelial systems at homeostasis and its response to inhaled allergens to enable mapping the downstream targets of tuft cells. *Alternaria* elicits strong innate and adaptive immune responses in the airways (67–70) initiated by the activation of several epithelial subsets including tuft cells, which respond by generating CysLTs (32). We performed scRNA-seq at homeostasis (**Fig. 3A,** left) and after nasal inhalation of the mold allergen *Alternaria alternata* in the innate phase (36 hours after a single inhalation) and adaptive phase (36 hours after 2 inhalations given 6 days apart) (**Fig. S5A**). We included equal numbers of CD45^+^ immune cells and EpCAM^+^ cells expanding the gating strategy to include EpCAM high and intermediate/OSNs, and again used microdissection to sample cells from olfactory and respiratory mucosa (**Fig. S5B**). In total we profiled 50,448 high-quality cells after filtering (**Methods**). Unsupervised cluster analysis (**Methods**) detected 11 subtypes of myeloid (**Fig. 3B, S5C)** and 17 subtypes of lymphoid cells (**Fig. 3C, S5D**).

**Fig. 3.**
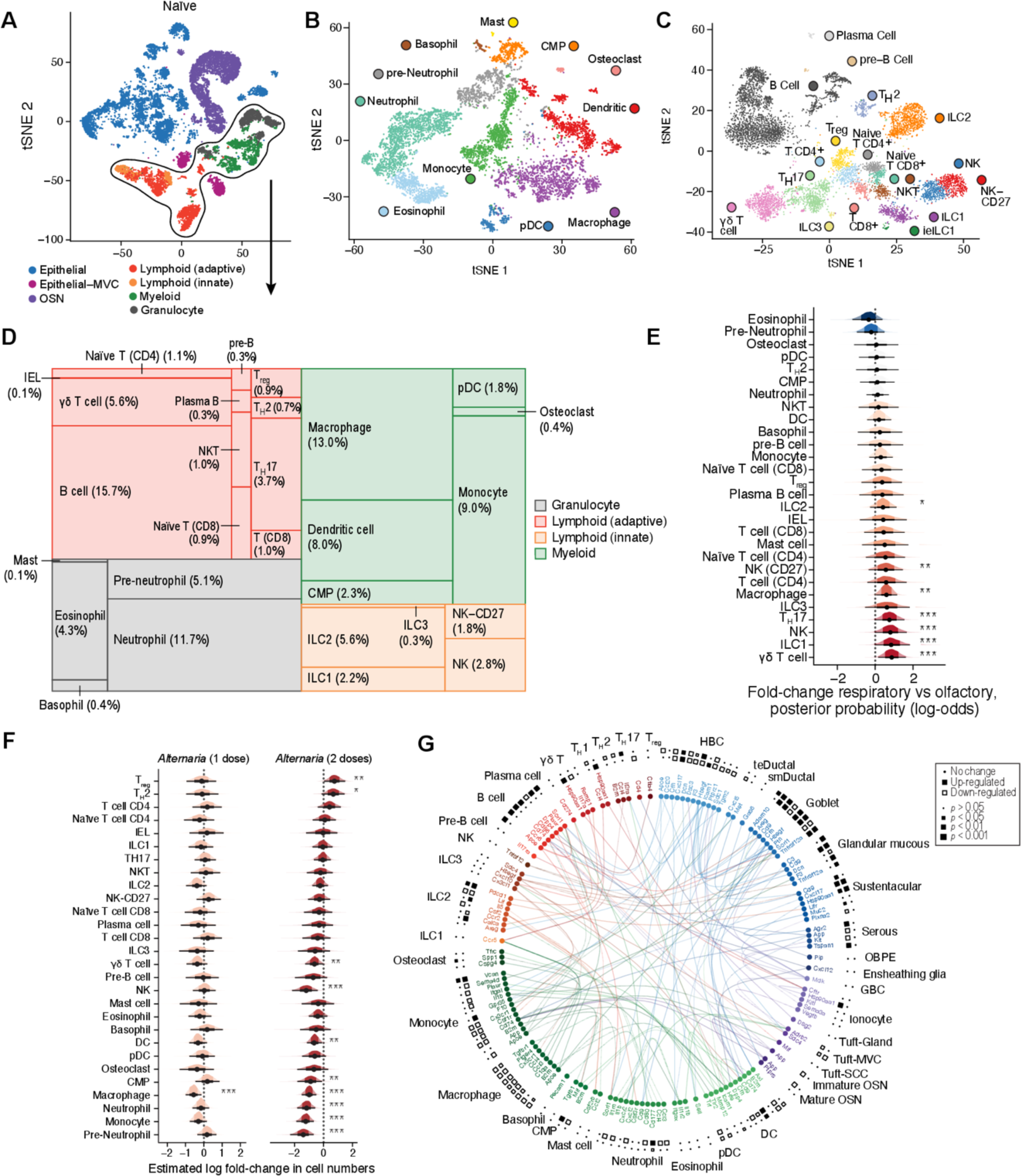
Single-cell analysis of intercellular communication and compositional shifts during allergic inflammation in the mouse nasal mucosa. (**A**) tSNE embedding of 19,275 cells from nasal mucosa of naïve mice (*n*=4 mice) assessed by scRNAseq, cell type lineages identified by unsupervised clustering are shown (color legend). (**B-C**) tSNE embeddings of 7,107 myeloid (**B**) and 7,213 lymphoid (**C**) cells (points). Cells are colored by the cell types (inset legend). (**D**) Tree map plot shows the cell-type composition (size of rectangles) of the immune compartment in naïve mice. **E-F**. Density histograms show the posterior distribution from a Bayesian Dirichlet Multinomial regression model (**Methods**) of log fold-change estimates (x axis) of the effect of tissue location **(E)** and for the effect of one (orange) and two (red) doses of *Alternaria* (F) on the proportion of each immune cell type. Black dot: point estimate, thick bar: 50% credible interval, thin bar: 90% credible interval (CI) * 90%, ** 95%, *** 99% CI does not include 0. (**G**) Circle plot displays differentially expressed receptor ligand interactions between cell types (color) after *Alternaria* treatment. Squares denote significance (size) of up (black) or down (white) regulation after one dose (inner ring) or two doses (outer ring) of *Alternaria*.

At homeostasis, in the myeloid compartment, the major clusters of cells were dendritic cells, macrophages and monocytes as expected (**Fig. 3B, D**), but we also observed a substantial fraction of tissue-resident neutrophils (11.7%), along with eosinophils (4.3%), basophils (0.4%), and mast cells (0.1%), osteoclasts (0.3%) and a *Kit^+^/Cd34^+^* subset which likely represents common myeloid progenitors (CMP, 4.9%). Notably, we found that granulocytes and their precursors were localized at higher concentration in the olfactory mucosa consistent with classical immunomorphologic studies and recent studies of neutrophil populations in the olfactory compartment (**Fig. 3D, S5E**) (71). In the lymphoid compartment, we identified all three previously described innate lymphoid cell (ILC) subsets. Again, consistent with the known predominance of the lymphoid compartment in the nasal associated lymphoid tissue (NALT) at the floor of the nasal cavity, we find that lymphocytes are concentrated in the nasal respiratory portion (**Fig. 3E**). The ILC2 subset accounted for 5 and 6% of the hematopoietic cell pool in the olfactory and respiratory epithelia respectively (**Fig. S5E-F, Table S9**). We recovered all canonical T helper subsets except for T_H_1 cells and observed a population of γδ T cells which expressed high levels of *Il17a* and *Rorc* (**Fig. S5D**) as did T_H_17 cells. Both *Il17a*-expressing subsets were significantly enriched in the respiratory epithelium (**Fig. S5D-F**), as were γδ T cells, ILC1, and NK cells (**Fig. S5D-F**).

### Inflammatory response to aeroallergen inhalation in the mouse nasal mucosa

*Alternaria* inhalation triggered rapid recruitment of eosinophils and ILC2 expansion in the nose (**Fig. S6A**), similar to the immune response in the lung when assessed by FACS (31, 32, 41, 67, 68, 70, 72). Analyzing far fewer cells, our single-cell data was not statistically powered to detect this change, and showed no significant differences (**Fig. 3F**). Interestingly, we also found a moderate increase in the recently described population of SiglecF^+^ olfactory-specific neutrophils (71). ScRNA-seq analysis of the immune compartment revealed relatively stable numbers of immune cell subsets the initial response 1 day after a single inhalation of *Alternaria* allergen, except for a 43.0% [CI-95: 12.3, 62.9] decrease in macrophage numbers (**Fig. 3F** and **S6A**). The discrepancy between FACS and scRNAseq approaches in identifying eosinophils is likely explained by their low mRNA content. During the adaptive phase of inflammation 36 hours after the second of 2 doses of *Alternaria*, we observed increased number of polarized T cells, particularly a 1.9-fold [CI-95: 0.87, 4.3] increase in activated T_H_2 cells as expected, but also a 2.1-fold [CI-95: 1.03, 4.42] increase in regulatory T (T_reg_) cells (**Fig. 3F**). Concomitant with the increase in adaptive lymphocytes, we observed a marked reduction in the fraction of almost all myeloid cell numbers (**Fig. 3F** and **Table S8**), likely a reflection of their stable numbers in the nasal mucosa.

Next, we defined cell-type specific genes (**Table S5**) and performed differential expression (DE) analysis (**Fig. S6C-E, Methods**). *Alternaria* drove a type 2-polarized innate immune response 24 hours after inhalation with type 2 effector cytokines *Il5* (FDR=0.007) and *Il13* (FDR=0.302) specifically upregulated by ILC2s (**Table S6**). Type 2 helper T (T_H_2) cells showed the strongest transcriptional response to a second *Alternaria* dose (**Fig. S6E)**. We additionally observed up-regulation of *Ltc4s* in dendritic cells, eosinophils and tuft-MVCs (all FDR<0.25, **Table S6**), consistent with increased production of CysLTs – including by tuft cells – as part of the early innate inflammatory response.

Finally, we characterized the intercellular signaling induced by the aeroallergen among the diverse cell types of the nasal mucosa (**Fig. 3G**). Following the approach we previously developed (73), we mapped all receptor-ligand gene pairs – annotated in either CellPhoneDB 2.0 (74) or the FANTOM5 (75) databases – which showed differential expression in response to *Alternaria* (**Methods, Table S7**). In addition to the classical cytokines *Il5* and *Il13*, additional effector signaling associated with type 2 inflammation was upregulated by ILC2s including *Calca,* (**Fig. 3G**) and *Il17rb* (**Table S6**). Consistent with this type 2 polarization, the activation receptor *Ilr2a* on T_H_2 cells was up regulated, while the NK and ILC1 chemoattractant *Ccl4* (associated with type 1 inflammation) was reduced, along with a marked reduction in *Il17a* expression by γδ T cells.

### Mapping the unique subtypes of epithelial cells in the mouse nasal mucosa

At homeostasis, OSNs have a limited lifespan and are replenished from GBCs, proliferating committed neuronal progenitors (76). OSNs are surrounded by several types of epithelial cells that support the structure of the neuroepithelium and ensure its proper function (4, 5). To define the transcriptional profile of the diverse epithelial cells (**Fig. 4A**) and their anatomical distribution across respiratory and olfactory mucosa (**Fig. 4B, S7A, S7F** and **Table S8**), we combined scRNAseq and immunolocalization studies. In addition to the ionocytes and tuft-MVCs we characterized earlier; we found several other specialized epithelial subsets with predominantly olfactory distribution. Consistent with the known organization of the olfactory and respiratory epithelium (76–78), our analysis identified two subsets of progenitor epithelial cells corresponding to HBCs and GBCs (**Fig. 1A, 1D**, and **S7A**). As expected, GBCs were highly enriched in the olfactory epithelium (**Fig. 4B**). Epithelial stem cells of the respiratory mucosa and the olfactory HBCs were transcriptionally highly similar despite their distinct morphology (*R*>0.95, **S7B-C**). We identified a few distinct markers including the respiratory-enriched *Krt15* (**Fig. S7D**) but refer to the respiratory epithelial stem cells as HBCs because of their high transcriptional similarities. Sustentacular and ductal cells, GBCs, tuft-MVCs and ionocytes were predominantly derived from the olfactory mucosa (**Fig. 4B**).

**Fig. 4.**
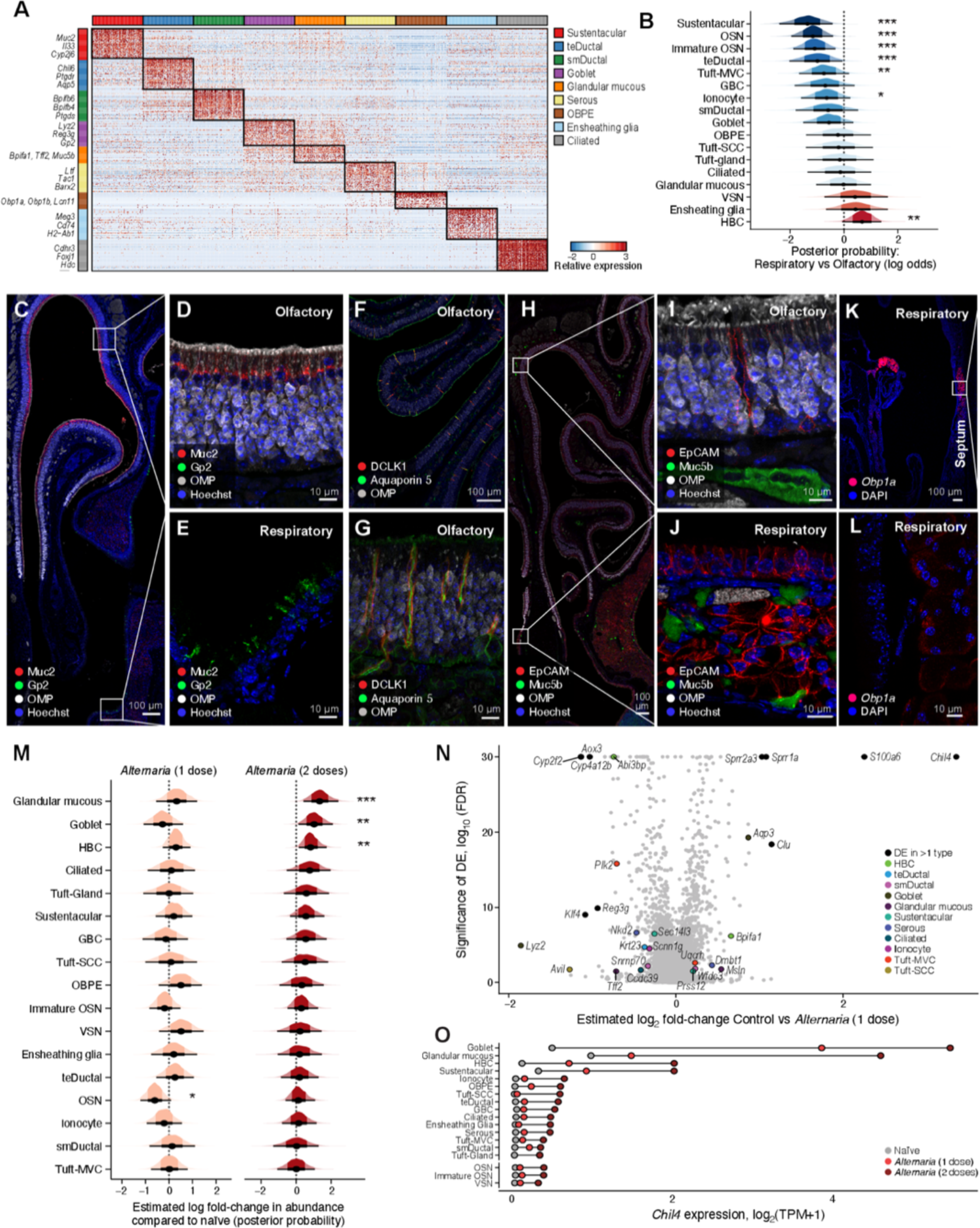
Mapping the unique subtypes of secretory epithelial cells in the nasal mucosa. (**A**) Heatmap shows the expression level (row-wise z-score of log_2_(TPM+1), color bar) of marker genes (right) specifically expressed (FDR<0.01) by each non-sensory epithelial subset (color bars). (**B**) Density histograms of log fold-change estimates (x axis) of effects of tissue location on proportions of each epithelial cell type. Black dot: point estimate, thick bar: 50% CI, thin bar: 90% CI. * 90%, ** 95%, *** 99% CI does not include 0. (**C-L**) Immunolocalization of epithelial cell subsets in the olfactory and respiratory nasal mucosa. (**C**) Cross section through the anterior section of the nose immunolabeled with OMP to delineate the olfactory epithelium in combination with Muc2 and GP2. Closeup of the olfactory epithelium (**D**, top inset of **C**) and respiratory epithelium (**E**, bottom inset of **C**) mucosa identifies the location of respiratory goblet cells marked by GP2 and sustentacular olfactory cells marked by Muc2. (**F)** Olfactory turbinates marked by OMP immunolabeled for DCLK1 in combination with Aquaporin 5. (**G**) Closeup of **F**. (**H**) Cross section through the posterior portion of the nose with olfactory turbinates (top), respiratory epithelium (bottom) and lateral nasal gland (lower right) demonstrating the distribution of Muc5b in the olfactory (labeled with OMP) and respiratory mucosa. (**I**) Closeup of olfactory epithelium and submucosa from **H**. (**J**) Closeup of respiratory mucosa of the nasal septum from **H** with epithelium overlying the submucosal glands. (**K**) Cross section through the anterior portion of the nose with septum on the right and lateral turbinates on the left demonstrating the expression of *Obp1a* mRNA detected by RNA scope ISH. (**L**) Closeup of *Obp1a* mRNA expression in the septal submucosal gland from K. (**M**) Density histograms of log fold-change estimates of changes in proportions of epithelial cell subsets after one (left) and two (right) doses of inhaled *Alternaria*. Black dot: point estimate, thick bar: 50% CI, thin bar: 90% CI. * 90%, ** 95%, *** 99% CI does not include 0. (**N**) Volcano plot shows the relationship between log_2_ fold change (x axis) and significance (y axis) of differential expression of genes (points) within epithelial subsets (color legend) after one dose of *Alternaria*. (**O**) Lollipop plot shows the expression level (x-axis) of chitinase-like protein 4 (*Chil4*), before and after inhalation of *Alternaria* (color legend).

Non-neuronal cells of the olfactory epithelium are classically identified by their expression of *Cyp2g1* (79) and *Krt18* (80). Multiple mouse models use *Cyp2g1* to specifically target the olfactory epithelium without an effect on OSNs (81, 82). Here, we identify two major subsets of *Cyp2g1* expressing cells with unique characteristics: in addition to sustentacular cells, which were marked by *Muc2* (**Fig. 4A**) a distinct second population of epithelial cells was positive for *Cyp2g1,* negative for *Muc2* and instead highly expressed markers of Bowman’s gland ductal cells including *Aqp4* and *Aqp5* (83) (**Fig. 4A, S7E**). We confirmed Muc2 was ubiquitously and highly expressed marker of sustentacular cells (**Fig. 4C-D**), and did not mark ductal cells in the olfactory mucosa (**Fig. 4C-E**). By reanalyzing published scRNA-seq data from human olfactory epithelium (OE) (54), we confirmed that *MUC2* is also specific to human sustentacular cells (*p*<0.0001, data not shown). Sustentacular cells were also the main site of *Il33* expression in the nasal mucosa, consistent with previous findings (84) (**Fig. 4A**).

Co-localization of Aquaporin 5 and DCKL1 defined the second subset of *Cyp2g1^+^* olfactory epithelial cells as the cells lining the transepithelial ducts of the Bowman’s glands (**Fig. 4F-G**). Notably, this specific subset of Aquaporin 5^+^ DCLK1^+^ cells was restricted to the olfactory neuroepithelium, and we refer to them as transepithelial ductal cells (**Fig. 4G**, **S7F**). The transepithelial ductal cells are enriched for expression of the PGD_2_ receptor DP2 (*Ptgdr*) (**Fig. 4A**), associated with mucus secretion in humans (85). In addition to the *Aqp5^+^/Dclk1^+^* transepithelial ductal cells, we identified a separate population of epithelial cells marked by very high expression of *Aqp5*, but negative for *Dclk1* and *Cyp2g1.* Immunofluorescent histology localized these cells to the submucosal portion of the Bowman’s gland ducts (**Fig. 4G**). Although rare, these submucosal ductal cells were quite distinct in their expression of several members of the bactericidal/permeability-increasing protein (BPI) fold family members – *Bpifb3, Bpifb4* and *Bpifb6* (**Fig. 4A**, **Table S5**). These genes initially named and defined for their sequence homology to *Bpifa1* (encoding the antimicrobial protein Splunc1), have now been shown to encode proteins associated with endoplasmic reticulum and Golgi vesicle trafficking, and their function has been associated with viral replication (86–88). This is notable, as the submucosal ductal cells are among the highest expressing subtypes for the SARS-CoV2 entry molecule ACE2 (**Fig. S7G-I**).

While Muc2 is restricted to the olfactory mucosa, we find two other subsets of secretory cells with broader distribution in the nasal mucosa: *Muc5b^+^/Tff2^+^* mucous cells and GP2*^+^* goblet cells. Our scRNA-seq data indicated that GP2^+^ goblet cells were more numerous in the olfactory portion while the glandular mucous cells were equally distributed in the olfactory and respiratory mucosa (**Fig. 4B**). Nasal goblet cells are marked by the expression of *Gp2* and genes encoding proteins with antimicrobial activity: *Lyz2* for lyzosyme C (89), and *Reg3g*, which encodes regenerating islet-derived protein 3-gamma, a bactericidal C-type lectin with activity against Gram^+^ bacteria (90) (**Fig. 4A**). Spatial analysis by immunofluorescence demonstrated that GP2 is widely expressed in luminal epithelial cells in the respiratory epithelium (**Fig. 4C**, **E**) and lateral nasal gland (**Fig. S7J-K**). The goblet cells of the lateral nasal gland, which is located in the posterior portion of the nose, account for all of the GP2 expression in the olfactory nasal portion (**Fig. 4C** and **Fig. S7J**).

The *Muc5b^+^* glandular mucous cells express the highest levels of several transcripts needed for epithelial defense in addition to their signature expression of *Muc5b*: the trefoil factor *Tff2,* secreted in concert with mucins and involved in epithelial restitution (35), as well as *Bpifa1* (Splunc1), important for nasal epithelial defense (91–93) (**Fig. 4A**). We used Muc5b protein as a marker of glandular mucous cells and found it was highly expressed in 1) the acinar cells of the submucosal Bowman’s glands of the olfactory epithelium (**Fig. 5H-I**); 2) the submucosal glands of the respiratory mucosa (**Fig. 4H, J**) and 3) the acinar cells of the lateral nasal glands (**Fig. 4H**). We detected low expression of Muc5b in GP2-expressing goblet cells lining the luminal surface of the respiratory epithelium and lateral nasal gland (**Fig. S7K-L**). Consistent with this wide distribution of mucin expressing cells in the nose, we find multiple Periodic Acid Schiff positive cells in the epithelial lining of the respiratory mucosa, septal and lateral nasal glands in addition to submucosal glands in the respiratory mucosa and in the Bowman’s glands of the olfactory mucosa (**Fig. S7M**).

**Fig. 5.**
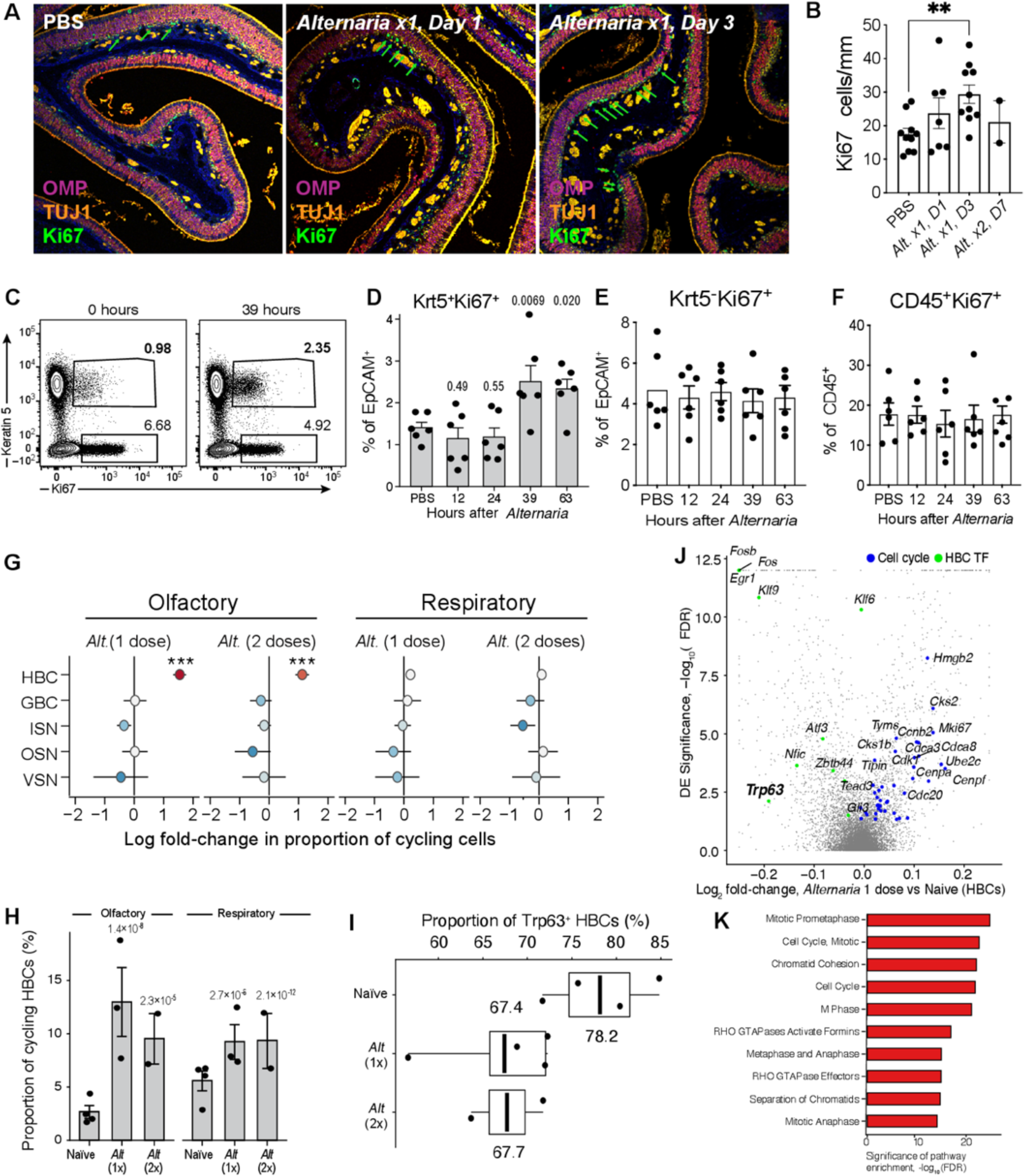
Induction of horizontal basal stem cell proliferation by allergen inhalation. (**A**) Histological assessment of the effect of *Alternaria* inhalation (right hand panels) on overall morphology, and on OSNs (OMP, purple), proliferating cells (Ki67, green) and ISNs (Tuj1, Orange) in the olfactory neuroepithelium compared to PBS controls (left). (**B**) Histological quantification of proliferating (Ki67^+^) cells at baseline and after *Alternaria*. (**C-F**) FACS analysis of Ki67 expression after a single *Alternaria* inhalation (30ug) at the indicated time points. Representative FACS plot demonstrating the expression of Ki67 among CD45^−^EpCAM^+^ cells before and 39 hours after *Alternaria* inhalation (**C**). (**D-F**) Quantitation of the percent of Keratin5^+^Ki67^+^ (**D**) and Keratin5^−^Ki67^+^ (**E**) among EpCAM^+^ cells and the proportion of Ki67^+^ cells among CD45^+^ immune cells (**F**). Data in **B, D-F** are means ± SEM from at least 2 independent experiments, each dot is a separate mouse, p-values: Wald test. **G-H.** Cell-type identity of *Alternaria*-induced proliferating cells from scRNA-seq data. **(G)** Estimated regression coefficients from Bayesian Dirichlet Multinomial regression (**Methods**) modeling the fold-change in proliferative state (x-axis) for each cell type (y-axis) after inhalation of *Alternaria* in the olfactory (left) and respiratory (right) nasal mucosa. Black dot: point estimate, thick bar: 50% CI, thin bar: 90% CI. • 90%, ** 95%, *** 99% CI does not include 0. **(H)** Estimated proportion (y-axis) of proliferating HBCs (Methods) in each mouse (dots) naïve mucosa and after inhalation of *Alternaria* (x-axis) assessed by scRNAseq. Bars show the mean and error bars show the SEM. **I-K**. Transcriptional activation of HBCs by *Alternaria* from scRNA-seq data. **(I)** Proportion of Trp63^+^ HBCs **(J)** Volcano plot shows differential expression (*y* axis shows -log10(FDR) and effect size (x axis) for HBCs after 1 dose of *Alternaria*. Cell-cycle and HBC-enriched TFs (FDR<0.001) are highlighted (color legend, top). **(K)** Statistical significance (x axis, -log_10_(FDR)) of top-ranked Reactome pathways enriched among up-regulated genes in HBCs.

Finally, we identified a unique population of epithelial cells marked by very high and specific expression of odorant binding protein (OBP)-encoding genes: *Obp1a, Obp1b, Obp2a* and other members of the lipocalin family: *Lcn11* and *Mup4* (**Fig. 4A**, **S7O, Table S5**). OBPs are a sub-class of lipocalins, defined by their property of reversibly binding volatile chemicals (odorants) for delivery to the olfactory receptor neurons (94, 95). Because of their high and unique expression of OBPs, we refer to these cells as OBP expressing (OBPE). Using *in situ* hybridization to localize OBPEs in the nose, we found clusters of cells with very high *Obp1a* mRNA expression consistent with the scRNAseq data (**Fig. S7N-O**). The *Obp1a* clusters were localized in the submucosal area of the lateral nasal gland similar to previous reports of expression of *Obp1a* in rats (94) (**Fig. S7N**). In addition, we found high *Obp1a* expression in the septal submucosal glands (**Fig. 4K-L**). Thus, we identified several sites that might contribute to the transport of odorants to OSNs.

After characterizing the homeostatic epithelial composition of the nose, we analyzed the differential responses of the olfactory and respiratory mucosa to aeroallergen exposure. We identified a 1.4-[CI-95: 0.6, 3.3] and 3.9-fold [CI-95: 1.5, 11.1] increase in glandular mucous cells in the epithelium after 1 and 2 *Alternaria* doses respectively (**Fig. 4M** and **Table S8**). We also found a 2.9-fold [1.1, 8.1] increase in GP2^+^ goblet cells after 2 *Alternaria* doses (**Fig. 4M**). We validated this increase in goblet cells using histology (**Fig. S8A-B**). Differential expression analysis (**Fig. 4N-O** and **S8C-D**) distinguished between cell-type specific and cell type-independent effects (**Methods**), which identified *Muc5b^+^* glandular mucous and *Gp2^+^* goblet cells as the most strongly affected cell-types (**Fig. S8C**). Finally, we identified chitinase-like protein 4 (*Chil4*) as the gene most strongly up-regulated after allergen challenge across multiple cell types (**Fig. 4N-O**). *Chil4* expression, previously shown to increase in lung type 2 inflammation (96) and in the nasal mucosa in an *Il13* overexpressing system (97) was differentially increased across many cell types after *Alternaria* treatment. Although the greatest increases of *Chil4* were in *Gp2^+^*goblet and *Muc5b^+^* mucous cells, notably *Chil4* expression increase was also significant in olfactory sustentacular cells and HBCs and even detectable in immature sensory neurons (ISNs) and OSNs (**Fig. 4O**). In addition to *Chil4, S100a6* and two small proline rich proteins associated with wound healing – *Sprr1a* and *Sprr2a3* and induced by olfactory neuroepithelial ablation (2) are differentially increased after *Alternaria* inhalation (**Fig. 4N**). In summary, our data indicates that the nasal epithelial response to *Alternaria* inhalation is highly plastic with induction of type 2 innate and adaptive immune responses and epithelial remodeling.

### Induction of horizontal basal stem cell proliferation by allergen inhalation

Aeroallergens such as *Alternaria* are known to activate epithelial cells in the respiratory mucosa leading to direct and indirect (immune cell-mediated) epithelial remodeling (31, 41, 68-70). Morphological studies have previously demonstrated that both acute and chronic inhalation of the mold *Aspergillus* are associated with inflammatory cell recruitment of eosinophils similar to what we found here **(Fig. S6A**) but also with rapid OSN and/or sustentacular cell death (98). To determine whether *Alternaria* causes similar disruption of the neuroepithelial structure, we first assessed the olfactory integrity in parallel with an assessment of the respiratory mucosa. The multilayered organization of the olfactory neuroepithelium with olfactory marker protein (OMP)^+^ mOSNs overlying 1-2 layers of β-tubulin^+^ (TUJ1^+^) ISNs and a single layer of proliferating Ki67^+^ stem cells was preserved after *Alternaria* (**Fig. 5A, S9A**). The thickness of the OMP layer did not change, implying there was no profound OSN loss (**Fig. 5A, S9B**). Assessment for olfactory apoptosis by the TUNEL assay demonstrated clear evidence of apoptotic cells in the respiratory – but not in the olfactory epithelium – after a single *Alternaria* inhalation (**Fig. S9C-D**). Confirming this finding, we found no induction by *Alternaria* of transcriptional pathways associated with necrosis or apoptosis in OSNs, while a minor signal was induced in HBCs and sustentacular cells (**Fig. S9E**). Consistent with this assessment, the proportion of OMP^+^ OSNs or sustentacular cells defined by scRNAseq did not significantly change (**Table S8**). Together these data conclusively show that *Alternaria* inhalation does not cause substantial cell death in the OE.

The most notable change in the olfactory mucosa was in the basal layer where the number of Ki67^+^ cells was visibly higher after *Alternaria* inhalation (**Fig. 5A**). Quantitation of Ki67^+^ cells in the olfactory epithelium indicated a significant induction of proliferation (**Fig. 5B**). In parallel assessments of the respiratory epithelium however, we noted that the number of Ki67^+^ respiratory cells was much lower at baseline, reflecting the lack of GBCs. Furthermore, the numbers of Ki67^+^ proliferating cells did not significantly increase after 1 or 2 *Alternaria* challenges (**Fig. S9F-G**) suggesting a profound effect of *Alternaria* on olfactory proliferation and a much more limited effect on nasal respiratory proliferation. As both GBCs and HBCs can act as stem and progenitor cells in the olfactory epithelium, we next sought to determine the identity of the olfactory cells induced to proliferate by the allergen. Immuno-fluorescence demonstrated that the Ki67^+^ cells in the olfactory epithelium are aligned at the basement membrane and partially co-localize with Keratin 5 (Krt5) suggesting they are HBCs (**Fig. S10A**). To further validate the identity of the proliferating cells using protein-level evidence, we employed an intracellular FACS assay for Krt5^+^ and Ki67. We confirmed that Krt5^+^ HBCs proliferate after *Alternaria* inhalation while the numbers of other proliferating immune and non-immune cells including Ki67^+^Kit^+^ GBCs did not change (**Fig. 5C-F** and **S10B-D**).

Finally, we leveraged our scRNA-seq data to define the activated cell populations at the transcriptional level. We examined the proportion of cycling cells of each type before and after *Alternaria* inhalation, finding that proliferation is strongly and specifically induced in HBCs (**Fig. 5G-H** and **S10E-F**). This effect was much stronger in the olfactory epithelium which showed a 6.1-fold increase (*p*=1.4×10^-8^, Wald test) in the fraction of *Krt5*^+^ cells which are positive for a cell-cycle gene signature (**Methods**) than in the respiratory (1.8-fold increase, *p*=2.7×10^-6^, Wald test) (**Fig. 5G-H**) consistent with the immunofluorescence assessment **(Fig. S9F-G)**. Proliferation of olfactory HBCs is induced by down-regulation of the transcription factor p63 (*Trp63*) (99), which our data confirms is specifically expressed by the HBC population (FDR<0.001, **Table S2**). At homeostasis, the majority (78.2%) of HBCs are *Trp63*^+^, compared to only 1.2% of non-HBC epithelial cells (**Fig. 5I**). We found that *Trp63* expression in HBCs was markedly reduced after a single dose of *Alternaria* (*p*=0.004) and also after two doses (*p*=0.02, **Fig. 5I-J**). Since p63 promotes olfactory stem cell self-renewal by inhibiting HBC differentiation (100, 101) and preceding HBC proliferation (99), this down-regulation is consistent with HBC activation by aeroallergen. *Alternaria* inhalation also induces down-regulation of the Notch signaling pathway (particularly *Notch1, Notch2* and *Jag2* all FDR<0.05, **Table S6**), consistent with known mechanisms of olfactory HBC activation (78, 102). Consistently, differential expression tests showed that up-regulated gene expression in HBC was dominated by cell-cycle related transcripts (**Fig. 5J**), and accordingly, the most significantly enriched pathways were all related to proliferation (**Fig. 5K**). Thus, we demonstrated that inhalation of the aeroallergen *Alternaria* induced proliferation of the quiescent HBCs in the 36-72 hours after exposure.

### Allergen-induced olfactory stem cell plasticity

Next, to define the plasticity of the olfactory stem cell niche in the setting of allergen exposure, we performed ‘pseudo time’ analysis on the 17,318 CD45^−^ epithelial and neuronal cells from the olfactory epithelium (**Fig. 6A** and **Fig. S11A-C**) using the partition-based graph abstraction (PAGA) algorithm (103). The inferred trajectory of cellular states (**Fig. 6A**) was consistent with the topology previously defined by Fletcher *et al.* (84). This demonstrated that HBCs give rise to GBCs, which in turn differentiate along a major pathway to OSNs, and a minor pathway to MVCs, which we show here to be tuft cells and ionocytes (**Fig. 1**). We identified the genes (**Fig. 6B, Fig. S11A-C** and **Table S9**) whose expression significantly varied (FDR<0.001, Wald test) with progression through neurogenesis (**Methods**). Early genes included canonical markers of HBCs, such as *Krt5* and *Trp63* (100), while genes at the latest stages included olfactory marker protein (*Omp*), and other markers of mature OSNs, such as *Chga* encoding chromogranin A*, S100a5* and *Olfm1* for olfactomedin 1 (**Fig. S11C**). As expected, GBC markers *Ascl1* and *Kit* were transiently activated during the intermediate stages of neurogenesis (**Fig. 6B**), while differentiation along with MVC lineage led to up-regulation of tuft cell markers *Trpm5, Ltc4s* and ionocyte markers *Foxi1, Ascl3* (**Fig. 6B**, bottom). Early in the fate branch toward OSNs, we observed up-regulation of a module of genes involved in regulation of cell cytoskeleton, including stathmin-1 (*Stmn1*), alpha (*Tuba1a, Tuba1b*) and beta-tubulin (*Tubb5*). Co-expression analysis of the scRNAseq data identified the subset of cells that are specifically induced by *Alternaria* inhalation as positive for a trio of markers *Mki67/Krt5/Stmn1* but not for *Mki67* and *Tubb3* (**Fig. 6C, S11**). This indicated that *Alternaria* induces a specific activation of the Krt5^+^ HBC stem cell pool and that the proliferative cells are marked by Stmn1^+^. We validated that stathmin-1 marks a subset of early precursors using immunofluorescence (**Fig. S12A**). By histology, very few proliferating cells were positive for tubulin 3 (TUJ1), a marker of mature and immature neurons (**Fig. S12B**), but some were positive for stathmin-1 (**Fig. 6D**).

**Fig. 6.**
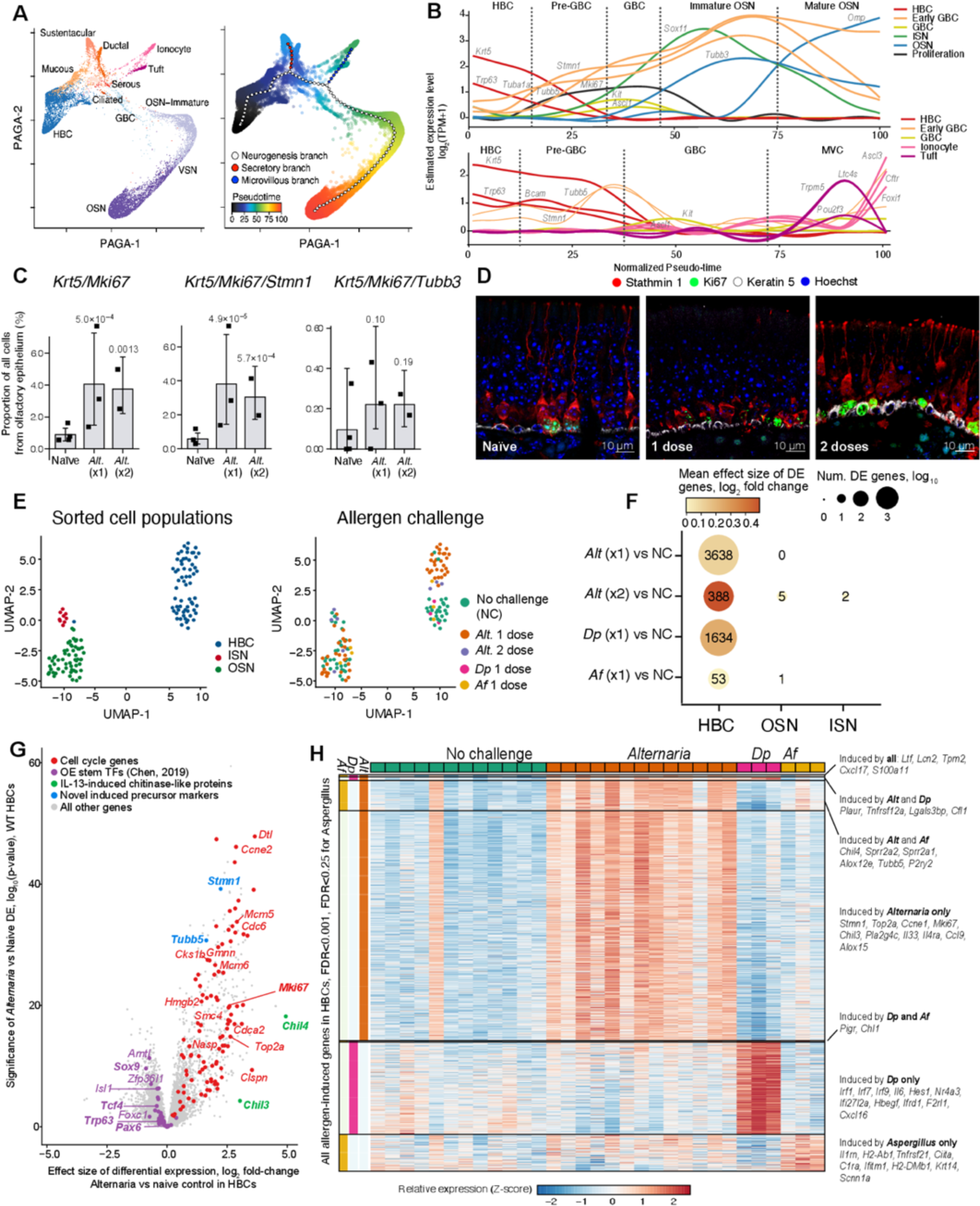
Mold and house dust mite allergens direct distinct olfactory stem cell activation programs. WT mice were given one or two inhalations of *Alternaria* and were assessed by scRNAseq (as in Fig. 3) (**A-C, G**) or by histology (**D**). (**A**) Scatter plots show low-dimensional PAGA embeddings of 17,318 olfactory epithelial cells (points) colored by cell-type cluster (left) and by pseudo-time coordinate (color legend, right). Trajectory (large points) and branches (color legend) were fit using elastic principal graphs. (**B**) Smoothed expression level (y-axis) of classical (*Krt5, Bcam, Trp63, Kit, Ascl1, Omp, Tubb3*) and novel (*Tub1a, Tubb5, Stmn1*) markers of intermediate stages of neurogenesis (top panel, color legend) and differentiation towards the microvillous cell fate (bottom panel) along the pseudo-time course (x-axis) from HBCs to mature OSNs. Stages (grey boxes) are shown as a visual guide. (**C**) Bar plots show the mean proportion of cells assessed by scRNAseq in which each individual (top row) and combination (bottom row) of marker genes is detected in each mouse (point). Error bars: 95% CI, p-values, likelihood-ratio test. (**D**) Protein expression of Stathmin 1, Ki67 and Keratin 5 in the olfactory epithelium of naïve mice (left) and after inhalation of one (center) and two (right) doses of *Alternaria* assessed by immunofluorescence. (**E-H**) WT mice were given a single inhalation of *Alternaria* (*Alt*), or *Aspergillus fumigatus* (*Af*) or *Dermatophagoides pteronyssinus* (*Dp*). Bulk RNAseq was performed on sorted HBCs (EpCAM^int^BCAM^+^), ISNs (EpCAM^int^NCAM1^+^) and OSNs (EpCAM^low/-^NCAM1^+^) a day after allergen challenge. (**E**) Uniform Manifold Approximation and Projection (UMAP) embedding of the bulk RNA-seq profiles of 130 FACS-sorted populations (color legend, left panel). **(F)** Dot plot shows the number (dot size, top-right legend) and mean fold-change (dot color, top left legend) of DE genes within each FACS-sorted population (x-axis) after each allergen challenge (y-axis). **(G)** Volcano plot shows the differentially expressed genes in HBCs after 1 dose of *Alternaria*, specific gene sets are labeled (color legend, top). **(H)** Heatmap shows the expression (row-wise Z-score, bottom color bar) in FACS-sorted BCAM^+^ HBCs from each mouse (columns) of 1373 genes (rows) up-regulated (FDR<0.001 for *Alternaria* and *Dp* and FDR<0.25 for *Aspergillus*) grouped into categories where they are induced (vertical color bars, left), selected genes are called out on the right.

Next, we developed a panel of cell surface antibodies to follow the dynamic response in the olfactory mucosa by FACS and high coverage bulk RNAseq of defined populations of HBCs, ISNs, and OSNs. Using a combination of three adhesion molecules – basal cell adhesion molecule (BCAM), a marker of stem cells (104), neural cell adhesion molecule 1 (NCAM1), a marker and epithelial cell adhesion molecule (EpCAM), we identified: HBCs (EpCAM^int^BCAM^+^), ISNs (EpCAM^int^NCAM1^+^), and OSNs (EpCAM^low^ NCAM1^+^) (**Fig. S12C**). In sorted cells, we confirmed that HBCs (EpCAM^int^BCAM^+^) express *Krt5*/Krt5 protein and *Trp63*, ISN (EpCAM^int^NCAM1^+^) express high levels of *Ncam1, Tubb3*/TUJ1 protein and the early neuron differentiation marker *Stmn1* while OSNs (EpCAM^low^NCAM1^+^) express the highest levels of *Omp*/OMP protein and low levels of *Stmn1* (**Fig. S12D-E**). Using these gates to isolate cells, we then used bulk RNAseq (**Fig. 6E**) to profile their transcriptomes after exposure to two separate mold allergens – *Alternaria* and *Aspergillus fumigatus* (*Af*) and the house dust mite allergen *Dermatophagoides pteronyssinus* (*Dp*) in WT mice. We found no dramatic changes in the proportion of HBCs, OSNs or ISNs after inhalation of any of those allergens (**Fig. S12F-G**). A single inhalation of each of those allergens caused significant differential expression in HBCs, but an additional *Alternaria* inhalation did not induce more profound changes (**Fig. 6F**). Consistent with our scRNAseq data, OSN and ISN gene expression profiles were not profoundly affected by *Alternaria* inhalation (**Fig. 6F**), and *Aspergillus* was similar (**Fig. 6F**). Up-regulated genes in HBCs of *Alternaria* challenged mice were prominently composed of cell cycle-related transcripts (**Fig. 6G**).

Along with *Trp63*, Fletcher (100) and Chen (82) identify three other TFs (*Pax6, Tcf4* and *Sox9*) necessary for maintaining HBC stemness, and all of these are down-regulated (FDR < 0.5) in HBCs by *Alternaria* (**Fig. 6G**). Indeed, genes down-regulated by *Alternaria* are enriched (*p*<0.01) for the complete set of 38 HBC stemness TFs (**Fig. 6G**) identified by Chen(82), providing further evidence of HBC activation, and the top up-regulated pathways were all cell cycle-related (**Fig. S13A**). Consistently (78, 102), Notch pathway signaling was also downregulated, including *Notch1, Notch2, Jag1*, and *Jag2* all FDR<0.0001 (**Table S10)**, and *Dll1* (FDR<0.05, **Table S10**).

We then compared the genes induced in HBCs by *Alternaria, Af* and *Dp*, to determine overlapping transcriptional programs and those specific to each allergen (**Fig. 6H**). The transcripts induced by all three allergens encode the monocyte-chemotactic chemokine *Cxcl17* and the antimicrobial peptides *Lcn2* and *Ltf.* Interestingly, *Cxcl17* and *Lcn2* are increased in the epithelium in the setting of allergic inflammation but shown to have a protective role with suppressing further inflammation (105). We found that the robust induction of an HBC proliferative response was unique to *Alternaria*, with induced stem cell activation markers *Mki67, Top2a* and *Stmn1* all up-regulated. In addition, *Alternaria* induced the chemokine *Ccl9,* the previously identified wound healing and IL-13 driven transcripts *Sprr2a1, Sprr2a2* and *Chil4* and *Chil3*. *Dp* inhalation was associated with a robust interferon response and induction of NFkB signaling but absent the robust proliferation response induced by *Alternaria* (**Fig. 6H, S13B**). Finally, *Af* inhalation led to a transcript induction of chitinase-like protein 4 (*Chil4*) expression to *Alternaria*, wound healing-associated transcripts *Sprr2a1* and *Sprr2a2* (106), and the arachidonate 12-lipoxygenase *Alox12e*, but with only a minor increase in HBC proliferation transcriptional programs (**Fig. S13C**). In summary, we find that inhalation of several allergens causes downstream activation of HBCs in the absence of profound effects on OSNs. While some pathways are shared, *Alternaria* prominently elicits the most profound transcriptional responses with induction of two distinct programs: stem cell proliferation and IL-13 dependent programs.

### Allergen-induced HBC proliferation depends on tuft cells, not type 2 inflammatory pathways

*Alternaria* inhalation triggers a potent innate immune response and downstream epithelial remodeling in the lung dependent on innate type 2 lymphoid cells and on STAT6 (69, 70). To understand whether the early olfactory proliferation and remodeling programs are dependent on innate lymphocytes and IL-13 signaling through STAT6, we administered a single *Alternaria* to *Il7r^−/−^*mice, which display reduced innate and adaptive lymphocytes (107) (**Fig S14B**), and in *Stat6^−/−^* mice, deficient in IL-4/IL-13 receptor signaling (108). Consistent with the findings in the lung, nasal inflammation (**Fig. S14A-C**), immune cell proliferation (**Fig. S14D**) and nasal eosinophil recruitment (**Fig. 7A**) were ablated in both *Il7r^−/−^* and in *Stat6^−/−^* mice. As ILC2-derived IL-13 is known to drive an epithelial stem cell proliferation and remodeling loop (34), we set out to determine whether this loop is responsible for the HBC proliferation in the nose. Surprisingly, we found that the proliferation of HBCs assessed by intracellular FACS of Keratin5^+^ cells was not reduced in either *Il7r^−/−^* or *Stat6^−/−^*mice (**Fig. 7B** and **S14E-F**). Consistent with the protein expression data, *Alternaria* induced *Mki67* expression equally in sorted BCAM^+^ HBCs from WT, *Stat6^−/−^* and *Il7r^−/−^* mice (**Fig. 7C, top**) as well as a previously validated gene signature of cell-cycle activation (**Fig. 7C, bottom**) (109). Finally, an *Alternaria-* signature score computed based on the expression of up-regulated genes from scRNAseq profiles of HBCs, was also unaffected in the *Stat6^−/−^* and *Il7r^−/−^*mice (**Fig. S14G**). We then sought to establish if other *Alternaria-*induced HBC transcriptional programs are dependent on IL-13 signaling and innate lymphocytes. Interestingly, the up-regulation of the IL-13 inducible gene *Chil4* (**Fig. 7D**) and the related chitinase gene *Chil3* (**Fig. S14H**) were completely abolished in both immune-deficient models. Overall proportions of HBCs, OSNs, and ISNs were unchanged by the genetic deletion of IL7R and STAT6 (**Fig. S14I**). Together these data establish that innate lymphocytes and STAT6 signaling are required for the induction of IL-13 directed HBC transcriptional programs but are dispensable for *Alternaria*-induced HBC proliferation.

**Fig. 7.**
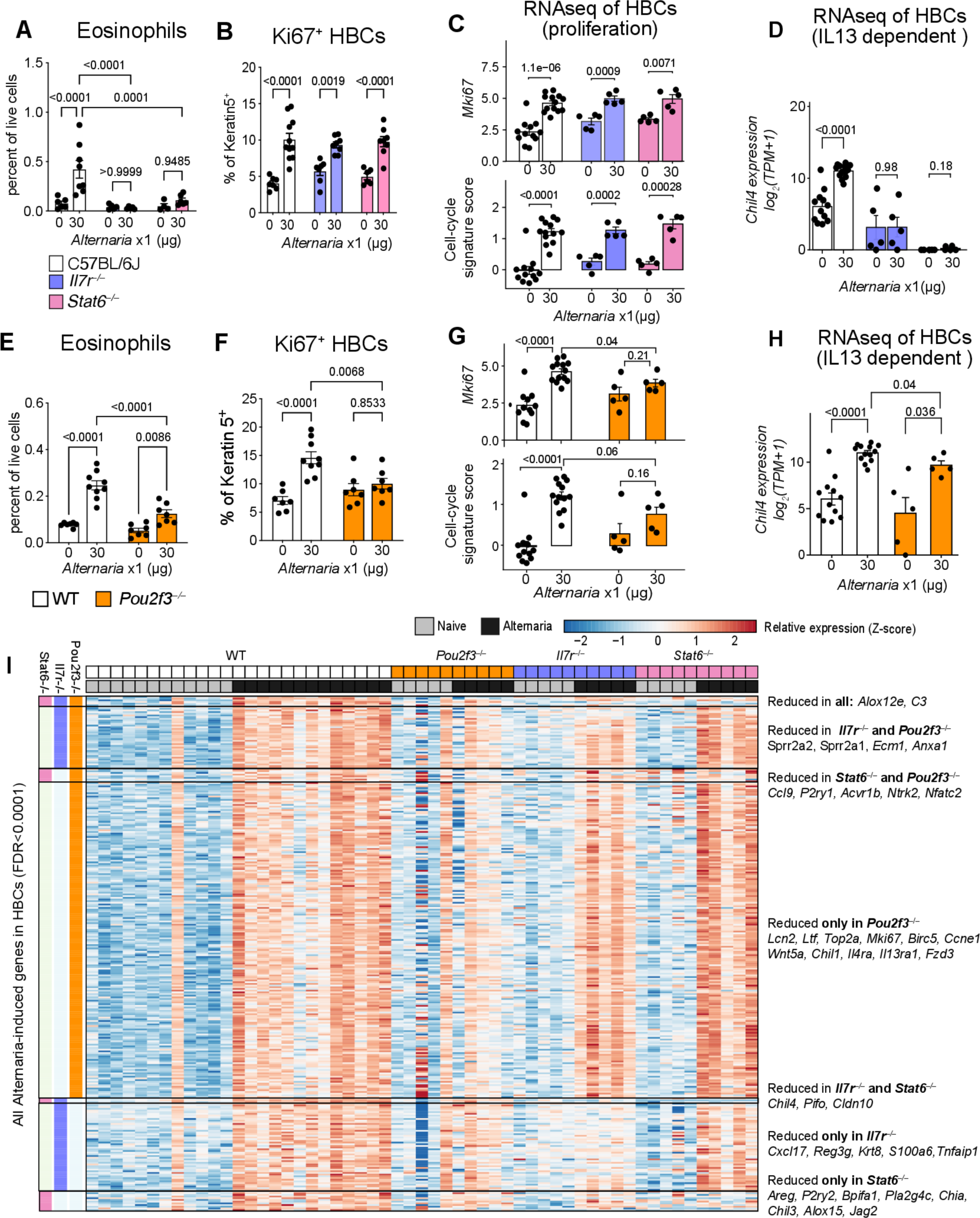
Allergen-induced olfactory stem cell proliferation is independent of type 2 inflammatory pathways and dependent on tuft cells. Mice of the indicated genotypes were given a single dose of *Alternaria* intranasally and assessed a day later. Number of eosinophils (**A, E**) and proportion of proliferating HBCs defined by Ki67 expression of Keratin 5^+^ cells (**B, F**) were assessed by FACS. HBCs were sorted based on BCAM and EpCAM expression (as in **Fig. 6, S12C**) and transcriptional changes were assessed by bulk RNA sequencing (**C-D, G, H**). Up-regulation of the cell cycle marker *Mki67* mRNA (top) and a previously published cell-cycle signature score (bottom, Methods) **(C, G)**, and similarly for the IL-13-dependent transcript *Chil4* (D, **H)**. **(I)** Heatmap shows the expression (row-wise Z-score, bottom color bar) in FACS-sorted BCAM^+^ HBCs from each mouse (columns) of 305 genes (rows) up-regulated (FDR<0.0001) by *Alternaria* and impaired (FDR<0.5) by at least one genotype (color legend), grouped into categories where they are impaired (vertical color bars, left), selected genes are called out on the right. Data are means ± SEM from three independent experiments, each dot is a separate mouse. Significance values in A, B, E and F are from one-way ANOVA with multiple comparison correction. All other p-values are Mann-Whitney U-test.

Tuft cells are directly activated by *Alternaria* (32), initiate lower airway inflammatory and remodeling programs some of which are independent of STAT6 signaling (31). To determine whether nasal tuft cells might regulate nasal inflammation, olfactory HBC proliferation and transcriptional programs through STAT6-dependent and independent mechanisms, we assessed the response to *Alternaria* in *Pou2f3^−/−^* mice that lack all mucosal tuft cells and specifically TRPM5^+^ tuft-MVCs (27) (**Fig. S4A-B**). Consistent with our previous findings in the lung (32) we found that eosinophil recruitment (**Fig. 7E**) was partially reduced in *Pou2f3^−/−^* mice, confirming the contribution of tuft cells to both upper and lower airway allergic inflammation. We also noted a higher baseline number of immune cells in the nose (**Fig. S15A**) which was mostly explained by an increased abundance of T lymphocytes (**Fig. S15B**) while homeostatic numbers of neutrophils and *Alternaria-*elicited neutrophil influx were largely unaffected by the absence of tuft cells (**Fig. S15C**). In addition, we noted a trend towards higher baseline of proliferating numbers of immune cells, which likely accounted for the lack of significant induction of immune cell proliferation in the *Alternaria-*challenged *Pou2f3^−/−^* mice (**Fig. S15D**). Thus, nasal tuft cells, similar to tuft cells in other mucosal compartments (110), are integral to nasal mucosal immune homeostasis.

Importantly*, Alternaria*-induced HBC proliferation was absent in tuft cell-deficient mice at the protein level assessed by intracellular FACS of Keratin5^+^ cells (**Fig. 7F** and **S15E-F**), as well as at the mRNA level (**Fig. 7G**, **top**). *Alternaria*-induced of the proliferation signature score described above were significantly decreased in tuft cell-deficient mice (**Fig. 7G**, **bottom**). Notably, this striking reduction of HBC proliferation in tuft cell deficient mice exceeded the magnitude of suppression of eosinophil recruitment. Consistent with the more modest effect on inflammation, we also found that the up-regulation of chitinase like protein transcripts *Chil3* (**Fig. S15H**), and *Chil4* (**Fig. 7H**) was only modestly affected in *Pou2f3^−/−^* mice. We then used differential expression tests to characterize the aspects of the *Alternaria*-induced transcriptional programs which depend on tuft cells (*Pou2f3*^−/−^), innate lymphocytes (in *Il7r*^−/−^) or IL-4/IL-13 receptor signaling (*Stat6*^−/−^). Of the 741 genes induced in HBCs by *Alternaria*, we found that 305 were impaired (negative fold-change and significant interaction term FDR<0.5) by at least one genotype (**Fig. 7I**). Of these, the majority (238, 78.0%) were impaired in tuft cell deficient mice, and of these, 114 were cell-cycle associated, including *Mki67*, demonstrating the dependency of the induced stem cell proliferative pathways on tuft cells. Conversely, we found that induction of *Chil4* was impaired in both *Il7r*^−/−^ and *Stat6*^−/−^ mice indicating its dependence on innate lymphocyte-derived IL-13. A further panel of IL-13 dependent transcripts were impaired only in *Stat6*^−/−^ mice including other chitinase-like proteins *Chil3, Chia*, and the 15-lipoxygenase *Alox15*(111), while the macrophage chemotactic *Cxcl17* and the asthma-associated transcript *Tmem45a* (112) were specifically impaired in *Il7r*^−/−^ mice (**Fig. 7I).** Together these data define the dependence of specific innate epithelial effectors molecules on the nasal mucosal immune system, and critically, define Pou2f3^+^ tuft cells as essential for the induction of olfactory HBC stem cell proliferation in response to aeroallergen.

## DISCUSSION

Tuft cells are solitary chemosensory epithelial cells scattered in most mucosal surfaces where they activate neural and immune circuits to promote inflammation and epithelial remodeling. The olfactory TRPM5^+^ MVCs share expression of the taste signaling channel TRPM5, the acetylcholine generating enzyme ChAT and the transcription factor Pou2f3 with the rest of the chemosensory family. They were previously considered distinct from the rest of the chemosensory tuft (also known as brush) family because of the low expression of taste receptors and lack of intimate connections with sensory neurons (47). Here, using microdissection of the nasal olfactory and respiratory mucosa, and single-cell analysis, we demonstrate that TRPM5^+^ MVCs share the core transcriptional profile of the chemosensory family and specifically its pro-inflammatory and remodeling mediator cassette. Furthermore, we identify a new population of glandular nasal tuft cells located in close proximity to odorant binding protein producing cells. We define the integrated olfactory remodeling and inflammatory response to the potent aeroallergen *Alternaria*. Unlike the dominant proinflammatory effect of *Alternaria* in the respiratory mucosa (69, 70), *Alternaria* induces proliferation of the quiescent HBCs. Allergen-induced HBC proliferation was dependent on TRPM5^+^ tuft MVCs, but independent of innate lymphocytes and STAT6 signaling. This establishes a distinct function of olfactory nasal tuft cells as specific drivers of stem cell proliferation.

Tuft cells in most mucosal compartments have been well characterized for their role in directing type 2 inflammation and downstream stem cell activation through an IL-13-driven tuft cell-ILC2 circuit (30, 34, 37). Here we found a more limited role for tuft cells as immune regulators, especially when compared to the degree of inflammatory cell reduction we observed in STAT6 deficient and lymphocyte depleted mice. Our data indicate that the dominant role of nasal TRPM5^+^ tuft-MVCs is the regulation of stem cell proliferation independent of inflammatory cascades. We considered several explanations for this discrepancy in tuft cell function in the olfactory vs other mucosal compartments: 1) distinct mediator profile, 2) unique multilayered organization of the olfactory neuroepithelium; and/or 3) the higher abundance of tuft cells in the nose.

In agreement with the previous literature, we found that TRPM5^+^ tuft-MVCs are distinct morphologically and in terms of receptor expression from the rest of the chemosensory airway tuft cell family (47). However, they are equipped with the same cytokine and enzymatic machinery that allows airway and intestinal tuft cells to direct immune responses – the acetylcholine generating enzyme *Chat*, eicosanoid cascade transcripts *Ptgs1* and *Hpgds* (for prostaglandins) and *Alox5, Alox5ap* and *Ltc4s* (for cysteinyl leukotrienes), and finally the transcript for the cytokine IL-25. We had previously shown that nasal ChAT-eGFP^+^ tuft cells generate cysteinyl leukotrienes and prostaglandin D_2_ upon allergen activation with *Alternaria* and *Dp* (32, 41). Now we conclusively demonstrate that >95% of these nasal tuft cells are TRPM5^+^ tuft-MVCs, establishing that these olfactory epithelial cells are directly activated by allergens and can generate pro-inflammatory mediators like the rest of the tuft cell family. Thus, an allergen can induce the classic tuft cell mediator response in the olfactory mucosa.

We then considered the possibility that the multilayered organization of the olfactory epithelium might provide a “buffer” of approximately 100µm between the apically positioned tuft cells and the submucosal immune system. Interestingly, the HBCs are also not in immediate contact with TRPM5^+^ MVCs, thus an intermediate cell might be activated here as well. Such a relay of TRPM5^+^ MVC cell derived acetylcholine signaling to IP3R3^+^ MVCs (ionocytes) was suggested by Lemons *et al*, who also found a moderate reduction in basal stem cell proliferation in *Pou2f3^−/−^* mice, albeit only after repeated inhalation of a mixture of odorants (113). Our more robust findings might be explained by the potent direct activation of TRPM5^+^ MVCs by allergens leading to CysLT production (32). LTC_4_, the biosynthetic product generated by tuft cells, is short lived and rapidly metabolized by extracellular enzymes (114). Its downstream products, especially LTE_4_ have a long half-life and are detectable in body fluids distant to the site of generation and thus could be the relevant downstream activators of stem cells. Consistent with this possibility, LTE_4_ inhalation is sufficient to induce an increase in tracheal tuft cells in a STAT6-independent manner, and its receptor is not directly expressed on tuft cells, suggesting that activation of an intermediate progenitor cell (31). This *in vivo* conversion of CysLT metabolites might account for the relay of tuft cell activation to stem cell proliferation suggested by Lemons *et al.* Interestingly, tuft cell deletion in the intestine is also associated with a significant reduction in the baseline number of proliferating epithelial cells (115). Thus, a role of tuft cells in directing proliferation of epithelial cells has been suggested before. However, in most mucosal compartments tuft cells are rare, accounting for approximately 0.5% of epithelial cells (116, 117), while olfactory tuft cells account for 6-7% of of EpCAM^+^ cells in the olfactory neuroepithelium. It is therefore likely that the combination of the abundance of tuft cells in the olfactory mucosa in combination with the unique organization of the olfactory neuroepithelium accounts for the dominant effect of olfactory TRPM5^+^ MVCs on HBC proliferation.

The olfactory neuroepithelium has at least two populations of basal cells (78). HBCs share many morphologic and transcriptional features with other mucosal epithelial stem cells. GBCs, unique to the olfactory epithelium, provide a pool of progenitor cells that repopulate OSNs and some differentiated epithelial cells (16, 76). At homeostasis HBCs are quiescent, while GBCs are continuously proliferating, but in the setting of injury and inflammation both HBCs and GBCs contribute to the regeneration of the olfactory neuroepithelium (2, 16, 81, 82, 102, 118). Thus, the specific activation of HBCs or GBCs might depend on the environmental signals that direct the mucosal response. OSN and sustentacular cell apoptosis are two potent inducers of both HBC and GBC proliferation (2, 16, 118) and so are sustained inflammatory signals such as TNFɑ and IL-13 (82, 97). Here we found an unexpectedly robust stem cell response without widespread cell death or changes to the overall architecture of olfactory neuroepithelium. Surprisingly, although we found clear evidence of a type 2 inflammatory response after allergen inhalation, the proliferation of stem cells was independent of the ILC2-mediated IL-13 signaling that directs stem cell activation in other mucosal compartments.

Using a combination of structural, proliferation analysis, and single-cell transcriptional mapping of the olfactory response to *Alternaria,* we identified the allergen-induced proliferating stem cell as quiescent HBCs (102, 118-120). Given the absence of pronounced necrosis or apoptosis, the *Alternaria*-induced activation we demonstrate here provides a physiological model of olfactory HBC activation. HBC activation is known to depend on the transcription factor p63 (*Trp63*), which promotes olfactory stem cell self-renewal by inhibiting differentiation (100, 101), and its down-regulation is required prior to HBC proliferation (99). Our data confirm previous findings that *Trp63* is specifically expressed by the HBC population (101), and our analysis of *Trp63* and Notch signaling using single-cell (**Fig. 5J**) and bulk (**Fig. 6G**) transcriptomics showed that along with HBC-specific down-regulation of *Trp63, Alternaria* inhalation also induced down-regulation of the Notch signaling pathway, both consistent with the mechanisms of olfactory HBC activation described by the Schwob group (78, 102).

Our analysis of *Alternaria-*exposed HBCs in *Pou2f3*^−/−^, *Il7r*^−/−^, and *Stat6*^−/−^ mice enabled us to dissect the specific aspects of the integrated response which are contributed by tuft cells, lymphocytes, and IL-13 signaling respectively. Perhaps surprisingly, we found that of the three, the loss of tuft cells had the most profound impact on the transcriptional response, perhaps explained by their position as early and direct sensors of allergens. Consistent with this possibility, we observed that the allergen-induced expression of the receptors *Il4ra* and *Il13ra1* was reduced in tuft cell deficient mice.

Several critical epithelial effectors involved in barrier defense, which are also all implicated in asthma, including the protective signaling proteins SPLUNC1 (*Bpifa1* (121, 122)) and Amphiregulin (*Areg* (123)), and anti-fungal chitinase-like proteins (*Chia, Chil3* (124)), were all impaired in *Stat6*^−/−^, but not *Il7r*^−/−^ or *Pou2f3*^−/−^, indicating their dependence on a non-lymphoid source of IL-13. Our single-cell profiling of the nasal immune compartment implicates mast cells and basophils as this potential source, as their expression of *Il13* is highest apart from ILC2s. Interestingly, the set of *Alternaria*-induced genes impaired in both *Stat6*^−/−^ and *Il7r*^−/−^ mice was extremely small, which suggests, perhaps surprisingly, that the direct regulation of HBCs by ILC2-derived IL-13 is a relatively minor pathway. On the other hand, the program impaired exclusively in *Il7r*^−/−^ was more extensive – indicating that lymphocytes, likely ILCs, regulate HBCs using IL-13-independent mechanisms – and included key modulators of the nasal immune response, such as the chemokine *Cxcl17*, recently implicated in macrophage-mediated airway remodeling (125).

Although we identified tuft cells as the critical link between allergen sensing and HBC proliferation, we did not delineate the downstream signaling pathways that link allergen activation of tuft cells and olfactory stem cell proliferation. Furthermore, we found no evidence that the proliferating basal cells then develop into OSNs or differentiated epithelial cells. Therefore, an important possibility is that tuft cells induce a proliferative state in basal cells that is not associated with regeneration, but is part of an *aberrant* response to allergen sensing by the epithelium. Importantly, BCAM^+^ stem cells (transcriptionally similar to the BCAM^+^ HBCs we assess here) were recently identified as the subset of basal cells that is specifically expanded in CRS with nasal polyps (104). These expanded stem cells account for the basal cell hyperplasia associated with CRS with nasal polyposis (51). As tuft cells were shown to both increase in numbers and assume an eicosanoid-enriched phenotype in CRS with nasal polyps (126), a tuft cell to basal cell proliferation link in human nasal disease is possible. Inhibition of IL-4/IL-13 receptor signaling with dupilumab as a treatment for patients with CRS with nasal polyps is clinically effective and rapidly reverses the type 2 inflammatory transcriptome (127). However, dupilumab is not disease modifying and its clinical effects are promptly undone upon discontinuation of treatment, with a rapid return of the underlying respiratory inflammation. Thus, a reprogramming of basal cells that is independent of IL-4/IL-13 receptor-mediated inflammatory signaling, as identified in this study, might be the reason that dupilumab is an effective treatment but not a long-term cure.

In sum, we provide extensive characterization of the rare subsets of nasal epithelial cells, conclusively identifying MVCs as transcriptionally belonging to the tuft cell and ionocyte families. We provide genetic evidence that olfactory tuft cells play a surprising role in regulating the proliferation of the stem cell compartment independent of classic inflammatory loops. This identifies a direct link between allergen sensing by tuft cells and activation of a reserve neural stem cell population. Together, these data deepen our understanding of the extensive epithelial heterogeneity in the nose and demonstrate that the functional capacity of tuft cells extends beyond triggering inflammation, as they are also potent regulators of tissue homeostasis and transformation.

## MATERIALS AND METHODS

### Study design

The aim of this study was to determine how olfactory TRPM5^+^ MVCs relate to the rest of the chemosensory family and define their function in the olfactory mucosa. We characterized the transcriptional profile of murine and human olfactory epithelial cells and the integrated response to allergen in the olfactory mucosa using machine learning approaches using three murine models of aeroallergen-induced innate inflammation (*Dermatophagoides pteronyssinus, Aspergillus fumigatus* and *Alternaria*). To delineate the contribution of tuft cells and tuft cell-initiated type 2 inflammatory cascades to *Alternaria*-elicited olfactory stem cell activation, we used the *Alternaria* allergen inhalation model in strains lacking tuft cells, innate and immune lymphocytes and deficient in IL-4/IL-13 receptor signaling. Mice were randomly assigned to treatment groups after matching for sex and age. All experimental replications are specifically described in the relevant figure legends.

### Mouse models

Mice were maintained at the Brigham and Women’s Hospital specific pathogen-free animal facility in accordance with the guidelines established by the Institutional Animal Care and Use Committee and Laboratory Animal Resource Center. The MGB Animal Care and Use Committee approved animal protocols in accordance with NIH guidelines. C57BL/6 wild type (WT) mice were originally purchased from Charles River Laboratories (Wilmington, MA), ChAT^BAC^-eGFP [B6.Cg-Tg(RP23–268 L19-EGFP)2Mik/J] (ChAT-eGFP) from Jackson Laboratories and *Pou2f3*^‒/‒^ described previously (27) were all bred in house. *Stat6*^‒/‒^ mice (B6.129S2(C)-*Stat6^tm1Gru^*/J), *Il7r*^‒/‒^ (B6.129S7-*Il7r^tm1Imx^*/J) and age and sex-matched WT C57BL/6J controls were purchased from Jackson Laboratories (Bar Harbor, ME). The mice used were between 11 and 36 weeks old. Pooled results include both male and female mice.

### Aeroallergen challenge protocols

For all *in vivo* allergen challenge experiments, mice were given intranasal inhalations of *Alternaria alternata* culture filtrate (lot #141774), *Dermatophagoides pteronyssinus* (lot# 49119) or *Aspergillus fumigatus* culture filtrate (lot# 195819) (all from Greer Laboratories) after sedation with an intraperitoneal injection of ketamine (10 mg/kg) and xylazine (20 mg/kg) and euthanized with ketamine overdose. The *Alternaria* culture filtrate was administered at a dose of 30 μg per mouse in 20 μl of sterile phosphate-buffered saline (PBS). The intranasal instillation was administered either as a single drop at the middle of the nose or as two separate administrations of 10 μl per nostril.

### Preparation of nasal cross sections

Snouts were collected from all mice after the mandibles and skin were removed. Snouts were immersed in 4% Paraformaldehyde (PFA) solution for 24 hours and then transferred to PBS. Fixed tissues were decalcified in 14% ethylenediaminetetraacetic acid (EDTA) for 14-24 days. Following decalcification, the nasal cavity was sectioned behind the incisors and in between the first three palatal ridges, yielding four coronal sections through the nasal cavity, and processed for routine paraffin embedding. Tissues were either frozen or embedded in paraffin. Paraffin embedded samples were sectioned to a thickness of 5 µm. Frozen samples were sections to thickness of 18 µm. Paraffin embedded sections were dewaxed in xylene for 20 minutes and hydrated through graded alcohols. The frozen sections were equilibrated to room temperature and post-fixed with methanol for 20 min.

### Immunofluorescence staining of nasal cross sections

Heat-induced antigen retrieval was performed in either Target Retrieval solution (pH 6) (Dako) or in EDTA (pH 8.5) (Sigma) for 30-40 min. Slides were then blocked and permeabilized with a PBS-based solution containing 0.1% Triton X-100, 0.1% saponin, 3% bovine serum albumin, and 3% normal donkey normal donkey serum for 1 hour at room temperature. All samples were stained with primary antibodies overnight at 4°C, rinsed with with PBS with 0.1% Triton-X (PSB-T) 3 times and incubated with secondary antibodies for 2 hours at room temperature unless otherwise indicated. The secondary antibodies included Alexa Fluor conjugates (488, 546, 594 and 647) (Invitrogen) (donkey anti-rat 594, donkey anti-mouse 594, donkey anti-mouse 488, donkey anti-goat 647, donkey anti-rabbit 546, donkey anti-rabbit 488, donkey anti-rabbit 647), donkey anti-chicken FITC (Invitrogen), all used at a concentration of 4 μg/ml. Nuclear staining was performed with Hoechst 33342 nuclear stain (Sigma).

The following primary antibodies were used after performing antigen retrieval with Target Retrieval solution (pH 6) (Dako): rabbit anti-DCLK1 (Abcam, 5 μg/ml), chicken anti-GFP (Abcam, 35 μg/ml), rabbit anti-neurogranin (Sigma Millipore, 70 μg/ml), goat anti-OMP (WAKO, 1 μg/ml), rabbit anti-gustducin (Santa Cruz, 1 μg/ml), rat anti-cochlin (Sigma Millipore, 10 μg/ml), rabbit anti-Stathmin1 (Invitrogen, 5 μg/ml), rat anti-GP2 (MBL, 2.5 μg/ml), rabbit anti-aquaporin 4 (Sigma Millipore, 5 μg/ml), rabbit anti-aquaporin 5 (Sigma Millipore, 2 μg/ml), rabbit anti-Muc2 (Abcam, 1.25 μg/ml), mouse anti-Muc5b (Abcam, 5 μg/ml), mouse anti-GP2 (MBL), rabbit anti-cytokeratin-5 (APC-conjugated, Abcam, 1.6 μg/ml), rat anti-Ki-67 (Biolegend, 2.5 μg/ml), rat anti-EpCAM (Biolegend, 2.5 μg/ml). For samples stained with rat anti-Ki-67 antibody (Biolegend, 2.5 μg/ml), the heat-induced antigen retrieval was performed in Target Retrieval solution (pH 6) (Dako) for 1 hour, prior to blocking as described above. Heat antigen retrieval for samples co-stained with rabbit anti-advillin (Novus Biologicals, 1 μg/ml) and chicken anti-GFP (Abcam, 35 μg/ml) was performed in EDTA (pH 8.5) (Sigma) for 40 minutes.

For double labeling of coronal sections with two primary antibodies derived from the same species (rabbit), the samples were incubated with the first primary antibody (neurogranin or Gɑ gustducin) overnight. The first rabbit antibody immunoreactivity was then developed with a donkey anti-rabbit fluorophore conjugated antibody. Residual rabbit immunoreactivity was subsequently blocked overnight with unlabeled donkey anti-rabbit Fab fragment (AffiniPure Fab Fragment Donkey Anti-Rabbit IgG (Jackson ImmunoResearch, 26 μg/ml). Tissues were rinsed with PBS-T and incubated with the second primary antibody raised in rabbit (DCLK1 or neurogranin). After PBS-T rinse, the immunoreactivity of the second rabbit antibody was detected with donkey anti-rabbit antibody conjugated to a different fluorophore.

### Goblet cell identification

Nasal tissue sections were also assessed by Periodic acid–Schiff (PAS) for identification of mucin-containing goblet cells. Slides were counterstained with hematoxylin for general morphologic examination.

### CFTR staining with tyramide boost

For CFTR detection in cross sections, target retrieval was performed with a citrate buffer (Target Retrieval solution, pH 6, (Dako) for 30min after deparaffinization. The samples were blocked for 1 hour with 10% goat serum, then incubated overnight with a mix of primary antibodies, containing rabbit anti-CFTR (Alomone, 16 μg/ml) and chicken anti-GFP (Abcam, 35 μg/ml) at 4C. The slides were then treated with poly-HRP conjugated secondary antibody from Tyramide SuperBoost™ kit (Invitrogen) and the developed with tyramide working solution with Alexa Fluor 594 from Tyramide SuperBoost™ kit (Invitrogen) for 7 minutes. The reaction was terminated with the stopping solution provided in the kit and subsequently incubated with goat anti-chicken Alexa Fluor 488 conjugated antibody (Invitrogen, 4 μg/ml) and Hoechst for nuclear stain for 2 hours.

Images were acquired at the Brigham and Women’s Hospital Confocal Microscopy Core Facility using the Zeiss LSM 800 with Airyscan confocal system on a Zeiss Axio Observer Z1 inverted microscope with 10× Zeiss [0.30 numerical aperture (NA)], 20× Zeiss (0.8 NA), and a 63× Zeiss oil (1.4 NA) objectives.

### Imaging based quantitation of cell numbers

For quantitation of cell counts in cross sections, 4-5 representative images of respiratory mucosa were acquired, each area of 0.2-1 mm^2^. For estimating the frequency of neurogranin^+^, Gɑ-gustducin^+^ and double positive cells, individual cells were manually counted using ImageJ software (National Institute of Health, Bethesda, MD) and the length of basal membrane was measured in microns. For neurogranin and ChAT-eGFP or Gɑ-gustducin co-staining assessment, the images were collected as 3-11 representative images of olfactory and 3-5 images of respiratory epithelium, each area of 0.8-1 mm^2^. To estimate the frequency of DCLK1-positive and neurogranin-positive cells in the lateral nasal gland (LNG), the length of basal membrane of LNG was measured in microns. The total number of ChAT-eGFP, neurogranin, Gɑ-gustducin, and Ki-67 immunoreactive cells in mouse nasal samples were manually counted using ImageJ software (National Institute of Health, Bethesda, MD).

For assessment of OMP thickness, 2-3 representative images of olfactory turbinates with total area of 0.28 mm^2^ were collected from ChAT-eGFP mice. To determine the thickness of olfactory mucosa, the area of the OMP^+^ layer was measured in mm^2^ using the ImageJ software.

### In situ hybridization (RNAscope)

The nasal snout for *in situ* hybridization was fixed in DEPC-treated 4% PFA and decalcified with DEPC-treated EDTA solution until sufficiently decalcified to section for OCT embedding. The tissue samples were sectioned at a thickness of 18 μms. For RNAscope *in situ* hybridization we used RNAscope® Fluorescent Multiplex Reagent Kit v2 (Cat No. 323120, Advanced Cell Diagnostics) assay following the manufactureŕs instructions. The nasal sections were post-fixed in 4% PFA for 15 min and then hydrogen peroxide for 10 min, both at room temperature. The sections were subsequently treated with Protease III for 10 min at 40°C before probe incubation. The probe to *Lct4s* was custom-made (Cat. No. 1092591-C1, Advanced Cell Diagnostics) and it targets exons 1-5 of the *Lct4s* transcript (nucleotides 98-534 of NM_008521.2). Expression of odorant binding protein 1a was evaluation using the mouse *Obp1a* probe (Cat. No. 500681, Advanced Cell Diagnostics). To mark tuft cells after *in situ* hybridization, sections were rinsed in PBS after final RNAscope wash and permeabilized and blocked again with 5% normal donkey serum/0.5% Triton X-100 Sorensońs buffer and incubated with a mix of antibodies to neurogranin and advillin in a dilution of 1:100 over night at 4°C. After several washes with PBS, sections were incubated with a Donkey anti rabbit antibody coupled to Alexa Fluor 647 for 2 hours and mounted in DAPI-FluoromountG. After RNAscope and/or immunostaining, the sections were imaged on a Zeiss Axio Observer Z1 inverted microscope. The entire sections were imaged in consecutive z-slices separated by 0.21 µm using a 63x oil objective. The z stacks were then projected at maximum fluorescence intensity using ImageJ.

### Whole septum imaging

For whole septum mount, the decalcified snouts were open longitudinally from the top to isolate the septum. The isolated septum was then permeabilized in the PBS-based solution described above for 3-4 hours. Primary chicken anti-GFP (Abcam, 35 μg/ml) antibody was added directly to the blocking buffer for incubation at 4°C for 72 to 96 hours. The specimens were then washed in PBS containing 0.1% Triton X-100 for 3 to 4 hours, and immunoreactivity to GFP was detected with secondary donkey anti-chicken secondary antibody, FITC (Invitrogen), applied for 72 hours at 4°C. Nuclear staining was performed with Hoechst 33342 nuclear stain (Sigma). The nasal septum was embedded using a glycerol-based mounting media and hardened under applied pressure, and 3-4 photographs of olfactory and respiratory epithelium, corresponding to 0.02-0.03 mm^2^ of total area per septum, were obtained. The surface area of individual tuft cells was measured using ImageJ software (National Institute of Health, Bethesda, MD) and averaged per biological replicate. To calculate the frequency of tuft cells, the number of GFP^+^ cells per image (ranging from 12 to 24 cells) was divided by the area captured in the focal plane.

### Human sample immunohistochemistry

Subjects between the ages of 18 and 75 years were recruited from the Brigham and Women’s Hospital (Boston, Mass) allergy and immunology clinics and otolaryngology clinics. The local institutional review board approved the study, and all subjects provided written informed consent. Ethmoid sinus tissue, middle or superior turbinate tissue was collected at the time of elective endoscopic sinus surgery from patients undergoing surgery for chronic rhinosinusitis or resection of pituitary adenoma. The samples were immediately placed in in RPMI medium (Corning, Corning, NY) with 10% FBS (ThermoFisher, Waltham, Mass) and 1 U/mL penicillin-streptomycin for transport to the laboratory on ice.

The samples were then fixed in 4% PFA for 4-8 hours and frozen in OCT for cryosectioning. The sections (6 μm) were post-fixed in methanol/acetone (1:1) mix for 15 min on ice. Heat-induced epitope retrieval was performed for 15 min in Target Retrieval solution (pH 6) (Dako) and all other steps were as described above for mouse snout processing. The primary antibodies included rabbit anti-advillin (Novus Biologicals, 1 μg/ml), mouse anti-NCAM (PE-conjugated, Biolegend, 0.4 μg/ml), goat anti-OMP (WAKO, 1 μg/ml), mouse anti-EpCAM (Santa Cruz,1 μg/ml) and rabbit anti-FOXI1 (ThermoFisher,0.5 μg/ml). The immunoreactivity was developed as described above with secondary antibodies raised in donkey.

### Single-cell suspension preparation for FACS, ex vivo stimulation and RNA seq

To isolate respiratory epithelium from olfactory mucosa, the tissue overlying the proximal part of the nasal septal and lateral wall, together with vomeronasal organ, was separated from olfactory epithelium, covering the distal portions of nasal cavity. The isolated olfactory and respiratory mucosa were processed as described previously (41). The separated tissue including cartilages and bone fragments was first incubated in a PBS solution with dispase (16 U/ml) and deoxyribonuclease I (DNase I; 20 μg/ml) for 30 min at room temperature on a shaker at 120 rpm. The digestion was terminated by adding cold DMEM supplemented with 5% fetal bovine serum (FBS). Then, the nasal mucosa was separated from the bones mechanically and incubated in Tyrode’s Solution with bicarbonate, HEPES, 0.25% BSA, without Ca^2+^ and added 31 U/ml of papain, 0.25 mg/ml L-cysteine and 20 μg/ml DNase I for 40 min at 37 °C on a rotating shaker at 220 rpm. The papain enzymatic activity was terminated with Tyrode’s Solution with HEPES and Ca^2+^ and with 0.01 mg/ml leupeptin and the digested mucosa was triturated thoroughly and prepared for flow cytometric staining. All samples were pre-incubated with TruStain FcX™ (Biolegend) at 1µg per 10^6^ cells in 100 µl volume for 10 minutes prior to immunostaining (**Fig. S1A**).

Single-cell suspensions from ChAT-eGFP mice were used for scRNAseq experiments at homoeostasis with specific enrichment of tuft cells (**Figs. 1-2**) and after *Alternaria* inhalation without enrichment of tuft cells (**Figs. 3-5**). All samples from ChAT-eGFP mice were stained with monoclonal antibodies against CD45 and EpCAM, both at 25 µg per 10^6^ cells in 100 µl volume for 40 min on ice. Propidium iodide (PI) was used as a dead cell exclusion marker and was added 1-3 minutes prior to sorting. To enrich for tuft cells, the epithelial cells from the respiratory and the olfactory mucosa were collected as three subsets of cells: globular tuft cells gated as EpCAM^high^ GFP^+^FSC^low^SSC^low^ cells, elongated tuft cells identified as EpCAM^high^ GFP^+^FSC^high^SSC^high^ cells, and other epithelial cells which were EpCAM^high^ GFP^-^ (**Fig. S1B-C**). The experiment was performed twice on two separate days each with 2 male and 2 female mice.

For assessment of the inflammatory and epithelial response to *Alternaria* by scRNAseq, the olfactory and respiratory epithelium were processed separately and CD45^+^ and EpCAM^+^ cells were sorted to represent 50% of cells in each location. The experiment was performed twice with the collected cells from olfactory mucosa of one mouse and respiratory mucosa from a different mouse of the opposite sex combined together to contribute equal proportions of a single sample. 6,000-10,000 cells per sample were collected into PBS and 0.04% (w/v) bovine serum albumin (BSA) and submitted for scRNA-seq. Single-cell suspensions for low input (bulk) RNA-sequencing were derived from the whole nasal mucosa. 1,000 cells per sample were sorted into TCL buffer (Qiagen) supplemented with 1% 2-mercaptoethanol.

For evaluation of HBC proliferation by intracellular staining, cell suspensions from the whole nasal mucosa were stained for extracellular markers with monoclonal antibodies against CD45, Kit (CD 117) and EpCAM, both at 25 µg per 10^6^ cells in 100 µl volume for 40 min. The samples were incubated for 1 hour with fixation buffer from True-Nuclear™ Transcription Factor Buffer Set (Biolegend) and washed with permeabilization buffer from the same set. The samples were then stained with intracellular markers Keratin-5 (Abcam, 3.2 µg/ml) and Ki-67 (Invitrogen, 20 µg/ml) for 45-50 minutes.

For evaluation of HBC, ISN and OSN transcriptional signatures, the single cell suspensions were incubated at room temperature for 40 min with antibodies against NCAM1, CD45, BCAM and EpCAM at 25 µg per 10^6^ cells in 100 µl volume. FACS sorting was performed using LSR Fortessa and 1,000 cells per sample were sorted into TCL buffer (Qiagen) supplemented with 1% 2-mercaptoethanol.

### Droplet-based single-cell RNA-Sequencing (scRNAseq)

Sorted samples, containing 6,000-10,000 cells each, were submitted for droplet-based single-cell RNA-sequencing (10X Genomics) to the Single Cell Genomic Core at Brigham and Women’s Hospital. Cells were washed and resuspended in PBS and 0.04% (w/v) bovine serum albumin (BSA) per the manufacturer’s guidelines. Purified samples were run through 10X controller and individually bar-coded libraries were generated and pooled. Pooled library samples were sequenced on a NovaSeq S1 (Illumina) in partnership with Dana Farber Cancer Institute (DFCI) Molecular Biology Core Facilities.

### Low input bulk RNA-sequencing

Cells from nasal single-cell suspensions were processed at the Broad Institute Technology Labs using low-input eukaryotic Smart-seq 2. Smart-seq2 libraries were sequenced on an Illumina NextSeq500 using a High Output kit to generate 2 × 25 bp reads (plus dual index reads). Computational processing of these data is described below.

## COMPUTATIONAL METHODS

### Pre-processing of scRNA-seq data

Demultiplexing, alignment to the mm10 transcriptome and UMI-collapsing were performed using the CellRanger toolkit (version 1.0.1, 10X Genomics). For each cell, we quantified the number of genes for which at least one read was mapped, and then excluded all cells with fewer than 500 or greater than 6000. We also excluded cells in which the fraction of transcripts mapping to the mitochondrial genome was greater than 0.25. Expression values *E_i_*_,*j*_ for gene *i* in cell *j* were calculated by dividing UMI count values for gene *i* by the sum of the UMI counts in cell *j*, to normalize for differences in coverage, and then multiplying by 10,000 to create TPM-like values, and finally calculating log_2_(TPM+1) values.

### Data partition

Our two single-cell datasets, the naïve nasal epithelium at homeostasis (**Fig. 1-2**) and after inhalation of *Alternaria* (**Fig. 3-6**), were analyzed separately. Additionally, because of the heterogeneity within the *Alternaria* dataset, we split this dataset by lineage into myeloid, lymphoid, and epithelial cells defined by unsupervised nearest-neighbor clustering (described below) and analyzed these subsets separately after identifying them through an initial analysis of the entire dataset. All downstream analyses described below, including final cell type identifications, were performed within these lineage datasets. The homeostasis dataset contained only epithelial cells and was processed in one group.

### Variable gene selection

For each gene, the relationship between sample detection fraction (cells in which at least one UMI was observed) and log of total number of UMIs was modeled using logistic regression. Outliers from this curve are expressed in a lower fraction of cells than would be expected, and are thus highly variable, that is, they are specific to a cell-type, treatment, condition or state. We selected these outliers using a threshold of deviance < -0.05 in all cases except for myeloid and lymphoid cell clustering where a threshold of -0.01 was used.

### Dimensionality reduction by PCA and t-SNE

We restricted the expression matrix to the subsets of variable genes and high-quality cells noted above, and values were centered and scaled before input to PCA, which was implemented using the R function ‘RunPCA’ included in the ‘Seurat’ package in R. For visualization purposes only (and not for clustering), dimensionality was further reduced using the Barnes–Hut approximate version of *t-*SNE. This was implemented using the ‘Rtsne’ function from the ‘Rtsne’ R package using 5,000 iterations and a perplexity setting of 50. Scores from the first 25 principal components were used as the input to *t-*SNE in *Alternaria* and homeostasis datasets, for the lymphoid and myeloid cells the first 50 PCs were used. After visualizing the data in this way, we performed cell clustering to identify cell types.

### Cell type identification using k-nearest-neighbor graph-based clustering

To cluster single cells by their expression profiles we constructed a Shared Nearest Neighbor (SNN) graph for our dataset using the built in Seurat function ‘FindNeighbors’. The number *k* of nearest neighbors was set at 20, and the SNN was constructed using the numbers of PCs described above. We used the SNN graph to identify clusters of cells using the Seurat ‘FindClusters’ function which implements the Louvain algorithm. We used a resolution parameter of 1.25 for the homeostasis and *Alternaria* datasets, we used resolutions of 1 and 3 for the myeloid and lymphoid subsets respectively. We then assigned cell type labels to these sets of cell clusters using known markers for nasal epithelial, immune, and neuronal subsets (**Table S1**). Because of the large proportion of certain cell types, they were frequently represented by multiple clusters, which expressed the same marker genes. Accordingly, we merged clusters expressing the same sets of known markers.

### Removal of doublets, technical artifacts, and contaminants

Clusters which co-expressed known markers for two distinct cell types, and which had elevated library complexity (number of detected genes) relative to those cell types, were considered doublets and removed. In this manner, 11 doublet clusters (3429 cells) in the *Alternaria* dataset and 8 clusters (586 cells) in the homeostasis dataset were removed. One additional cluster of 614 damaged or dying cells was excluded based on a particularly high proportion of mitochondrial reads, and marker genes consisting predominantly of mitochondrial genes. In addition, we removed two clusters were not consistently observed; that is, they were drawn predominantly from only a few biological replicates (mice). To determine this, we computed the number of cells within each cluster observed in each sample, and found that a linear regression described the mean-variance relationship in log-log space very well (adjusted *R*^2^=0.94), but that these two clusters were outliers (standardized residuals > 1.5) from this curve, indicating particularly inconsistent detection across mice, and were therefore removed. Finally, a cluster of 67 contaminating *Col1a1*^+^ fibroblasts was removed from the *Alternaria* dataset.

### Testing for differential expression

Differential expression (DE) tests were performed using MAST (128), which fits a hurdle model to the expression of each gene, consisting of logistic regression for the zero process (i.e., whether the gene is expressed) and linear regression for the continuous process (i.e., the expression level). All DE tests were run by comparing all cells of each type between conditions (downsampled to a maximum of 1,000 cells per condition). For each cell type, genes were only tested if they were detected in > 5 cells.

### Construction of cell-type signatures

To identify maximally specific genes for cell-types, we performed differential expression tests between each pair of clusters for all possible pairwise comparisons (downsampled to a maximum of 1,000 cells per group). Genes were tested if they were expressed in > 5 cells. Then, for a given cluster, putative signature genes were filtered using the maximum FDR *Q* value and the minimum log_2_ fold-change (across the comparisons). This is a stringent criterion because the minimum fold-change and maximum *Q* value represent the weakest effect size across all pairwise comparisons. Cell-type signature genes (**Table S2, S3, S6**) were obtained using maximum pairwise FDR=0.05 and a minimum log_2_ fold-change=0.1 in all cases except for lymphoid cell subtypes, where a more lenient cut-off of Fisher’s combined FDR=0.0001 and minimum log_2_ fold-change=0.2 were used.

### Scoring cells using signature gene sets

To obtain a score for a specific gene set (length *n*) in a given cell, a ‘background’ gene set was defined to control for differences in sequencing coverage and library complexity. The background gene set was selected for similarity to the genes of interest in terms of expression level. Specifically, the 10*n* nearest neighbors in the 2D space defined by mean expression and detection frequency across all cells were selected. The signature score for that cell was then defined as the mean expression of the *n* signature genes in that cell, minus the mean expression of the 10*n* background genes in that cell.

### Mapping nasal cell-types to tracheal using supervised machine learning

To compare nasal microvillous cells to recently characterized rare cell types in the trachea, we trained a random forest classifier using previously published single-cell profiles of tracheal epithelial cell types (36). The random forest was implemented using the “randomForest” R package. For training data we randomly sampled 250 cells per cell type, and used the 2000 most variable genes, the first 100 principal components, and 3 quality control metrics (number of genes detected, number of UMIs, and percent mitochondrial reads) as features. The model was then used to classify cells from the nasal dataset. Since some cell types would be exclusive to either nasal or tracheal epithelium and thus would not be expected to be correctly classified, we excluded low confidence predictions made by the model (with a classification probability < 0.5, computed as proportion of votes). A second random forest was fit in the same manner, this time using subtype-level classification of tuft cells (36), in order to identify subtypes of nasal microvillous cells.

### Identification of human tuft cell subsets

In the Durante (54) dataset the authors defined a subset of “Olfactory Microvillar Cells”. We restricted our analysis to this subset and reanalyzed these cells to compare them to the nasal microvillar cells in the mouse. We identified variable genes and clustered cells using the methods described above and used a resolution parameter of 0.25 for the final clustering which was then used for downstream analysis. Identification of DE marker genes was done using methods described above. Genes were tested if they were expressed in > 1 cell. For the Ordovas-Montañes dataset (51), no cluster of microvillar cells had been previously identified. We identified variable genes using a residual threshold of -0.001 and performed subsequent clustering and dimensionality reduction analysis using 50 principal components. Using a resolution of 3, we identified a cluster of 92 cells expressing known microvillous cell markers (*Foxi1, Cftr, Trpm5*). We then subset to this group and re-clustered it, identifying variable genes with a threshold of -0.01 and 5 principal components. Using a resolution of 1, we identified 6 clusters. One cluster expressed known tuft cell markers, and the remaining 5 expressed known ionocyte markers. We merged the Ionocyte expressing clusters and calculated marker genes with these two groups.

### Estimation of proportion of proliferating cells

All cells were scored for a previously defined cell-cycle signature gene set defined from scRNA-seq data (109) as described above. To define a data driven threshold for cycling vs non-cycling cells, the distribution of scores was clustered into two groups using an unsupervised mixture of gaussians implemented using the Mclust package in R (129). The proportion of the cells in the high-expression cluster was reported as the proportion of proliferating cells.

### Pseudo-time analysis of olfactory neurogenesis

We isolated the 17,318 CD45^−^ epithelial and neuronal cells (**Fig. 6A**, left) to examine the progression through neurogenesis using scanpy (130). The Partition-Based Graph Abstraction (PAGA) algorithm (103) was used to project cells into a low dimensional manifold, after defining unsupervised clusters generated using the Leiden algorithm (131). Elastic principal graphs (132) were then used to fit a branching tree through the PAGA co-ordinate space.

Spurious single-node branches were removed, producing a tree with a main branch from HBCs to mature OSNs, and two branches which were interpreted as a secretory and microvillous cell branch (**Fig. 6A**, right), consistent with earlier lineage tracing data (84). The node within the HBC cluster was manually selected as the root node for pseudo-time calculation. Genes significantly varying along the pseudo-time path we identified by fitting generalized additive models (133) to the expression level of each gene (**Fig. S11B-C**). Genes were ranked by the pseudo-time coordinate of their first non-zero fitted value in order to identify early markers of neurogenesis (**Fig. S11C**).

### Analysis of intercellular signaling networks

To analyze cell-cell signaling in our dataset, we looked for the presence of a set of interactions from the combined CellPhoneDB (74) and FANTOM5 (75) interaction databases. In order to ensure that the interactions in these sets represented feasible cell-cell interactions, we used the Jensen COMPARTMENTS database (134), and subset our interaction table to just those interactions in which one interacting partner was extracellular with high confidence (> 4.5). For plotting purposes (**Fig. 3G**), we identified receptors or ligands differentially expressed after *Alternaria* treatment (FDR < 0.05 and absolute log_2_ fold-change > 0.1). We used the 5 most significantly differentially expressed genes per cell type. We excluded genes with many interacting partner genes (≥10) to improve the visualization. For each differentially expressed gene in an interaction, its interacting partner was displayed in the cell type in which it was most highly expressed. Circle plot visualization was produced using the ‘edgebundleR’ R package.

### Bulk RNAseq pre-processing

Computational pipelines for RNA seq analysis were implemented as described elsewhere (35, 36). Briefly, sequencer generated BCL files were converted to merged, de-multiplexed FASTQ files using the Illumina Bcl2Fastq software pack-age v.2.17.1.14. Paired-end reads were mapped to the UCSC mm10 mouse transcriptome using Bowtie57 with parameters ‘-q–phred33-quals -n 1 -e 99999999 -l 25 -I 1 -X 2000 -a -m 15 -S -p 6’, which allows alignment of sequences with one mismatch. Gene expression levels were quantified as transcript-per-million (TPM) values by RSEM58 v.1.2.3 in paired-end mode (135). Expected count values were used for differential expression analysis using the DESeq2 R package.

### Bulk RNAseq quality control

Outlier analysis was performed using linear regression, fitted to an *Alternaria*-inducible gene expression score (top 20 differentially expressed genes in our scRNA-seq data). This procedure identified one mouse replicate which had a ‘difference in fits’ (DFFITS > 0.75), which was excluded from further analysis.

### Deconvolution of cell type composition within bulk RNAseq profiles

We determined the fraction of different cell types in each sample using CIBERSORTx (65) using default parameter settings. We provided our normalized log_2_(TPM+1) scRNA-seq data in (downsampled to a maximum of 200 cells per type) along with cell type cluster labels as input to CIBERSORTx.

### Testing for differences in cell proportions in bulk sequencing data

To model the compositionality of cell type proportions either within single-cell data or CIBERSORTx estimates within bulk sequenced cell populations, a Bayesian Dirichlet Multinomial Mixture (DMM) model was fit using the “brms” package in R (136). In the *Alternaria* dataset, cells from the epithelial and neuronal (EpCAM^+^) and immune (CD45^+^) compartments were collected separately. Cell proportions were therefore estimated as a fraction of their given compartment. The point estimates and posterior distributions for coefficients along with 50% and 90% credible intervals were then visualized as violin plots (**Fig. 3E, 4B, 4M**).

## Acknowledgments

We thank Adam Chicoine, Gerald Watts, Zhu Zhu and Kevin Wei from the Brigham and Women’s Hospital Center for Cellular Profiling for their invaluable assistance with cell isolation and single-cell RNA sequencing. We gratefully acknowledge Leslie Gaffney for help with arranging figures, and Daniel Montoro for technical advice. We are grateful to Charles Vidoudez and Sunia Trauger from the Harvard Center for Mass Spectrometry for their expertise and help setting up the mass spectrometry assays for CysLTs. We are also thankful to Dr. Lisa Goodrich at the Department of Neurobiology at Harvard Medical School for providing reagents and infrastructure for in situ hybridization. Schematic diagrams were created using BioRender.

## Author contributions

Conceptualization: ALH, LGB. Methodology: LGB, ALH, EL, SU. Investigation: SU, EL, LGB, ALH, AAB, CW, JL, DM, ECA. Writing: LGB, ALH, JAB, NAB, IM, EL, SU. Funding acquisition: LGB, JAB, NAB, ALH. Resources: TML, KMB, RR, AM, IM and JAB.

## Data and materials availability

All data is deposited in GEO (GSEXXXX) and in the Single Cell Portal (https://portals.broadinstitute.org/single_cell). All code required to reproduce the main steps of the analysis will be made available on request.

## List of Supplementary Materials

**Materials and Methods**

**Supplementary Figures:**

Figure **S1**: Isolation of single-cell suspensions from nasal respiratory and olfactory epithelium.

Figure **S2**: Anatomical and molecular characterization of OSN subtypes.

Figure **S3**: Three morphologically distinct types of nasal tuft cells share a common molecular profile.

Figure **S4**: Neurogranin acts as a novel tuft cell marker in multiple subtypes. Figure S5: Single-cell mapping of immune cell subtypes in the naive nasal mucosa.

Figure **S6**: Effects of allergic inflammation on gene expression and cell type proportions in the nasal immune system.

Figure **S7**: Expression of SARS-CoV-2 receptor ACE2 in epithelial cells and marker genes of differentiated mucosal subtypes in naïve mice.

Figure **S8**: Effects of allergic inflammation on gene expression and cell type proportions in the nasal epithelium.

Figure **S9**: Alternaria induces HBC proliferation in the absence of profound OSN damage.

Figure **S10**: Nasal olfactory mucosa architecture is preserved after Alternaria inhalation.

Figure **S11**: Pseudo-time analysis identifies novel marker genes for stages of olfactory neurogenesis.

Figure **S12**: Pseudo-time analysis identifies novel marker genes for stages of olfactory neurogenesis.

Figure **S13**: Transcriptional programs induced in HBCs by multiple aeroallergens.

Figure **S14**: Allergen-induced HBC proliferation is independent of type 2 inflammatory pathways.

Figure **S15**: Tuft cells are required for allergen-induced HBC proliferation.

## Supplementary Tables

Table **S1**: Known marker gene sets used for interpretation of cell clusters.

Table **S2**. Molecular signatures for all cell types in nasal epithelium.

Table **S3**. Molecular signatures for novel subsets of nasal tuft cells.

Table **S4**. Marker genes for human nasal tuft cells and ionocytes.

Table **S5**. Molecular signatures for epithelial and immune cell subsets in the nose.

Table **S6**. Cell type-specific gene expression dynamics after allergic inflammation.

Table **S7**. Modulation of cell-cell signaling during allergic inflammation.

Table **S8**. Cell type composition dynamics during allergic inflammation.

Table **S9**. Gene expression dynamics during olfactory neurogenesis.

Table **S10**. Allergen-induced inflammation dependent and independent transcriptional programs in the olfactory epithelium.

## References

1. D. B. Herrick, Z. Guo, W. Jang, N. Schnittke, J. E. Schwob, Canonical Notch Signaling Directs the Fate of Differentiating Neurocompetent Progenitors in the Mammalian Olfactory Epithelium. J Neurosci 38, 5022–5037 (2018).

2. L. Gadye, D. Das, M. A. Sanchez, K. Street, A. Baudhuin, A. Wagner, M. B. Cole, Y. G. Choi, N. Yosef, E. Purdom, S. Dudoit, D. Risso, J. Ngai, R. B. Fletcher, Injury Activates Transient Olfactory Stem Cell States with Diverse Lineage Capacities. Stem Cell 21, 775--790.e779 (2017).

3. C. Bachert, O. Krysko, P. Gevaert, M. Berings, C. Perez-Novo, K. van Crombruggen, in Mucosal Immunology, Fourth Edition. (2016), chap. 100, pp. 1899–1921.

4. T. Mao, B. Israelow, M. A. Pena-Hernandez, A. Suberi, L. Zhou, S. Luyten, M. Reschke, H. Dong, R. J. Homer, W. M. Saltzman, A. Iwasaki, Unadjuvanted intranasal spike vaccine elicits protective mucosal immunity against sarbecoviruses. Science 378, eabo2523 (2022).

5. J. E. Oh, E. Song, M. Moriyama, P. Wong, S. Zhang, R. Jiang, S. Strohmeier, S. H. Kleinstein, F. Krammer, A. Iwasaki, Intranasal priming induces local lung-resident B cell populations that secrete protective mucosal antiviral IgA. Sci Immunol 6, eabj5129 (2021).

6. S. A. Wellford, A. P. Moseman, K. Dao, K. E. Wright, A. Chen, J. E. Plevin, T. C. Liao, N. Mehta, E. A. Moseman, Mucosal plasma cells are required to protect the upper airway and brain from infection. Immunity 55, 2118–2134 e2116 (2022).

7. R. J. Hewitt, C. M. Lloyd, Regulation of immune responses by the airway epithelial cell landscape. Nat Rev Immunol 21, 347–362 (2021).

8. M. Zazhytska, A. Kodra, D. A. Hoagland, J. Frere, J. F. Fullard, H. Shayya, N. G. McArthur, R. Moeller, S. Uhl, A. D. Omer, M. E. Gottesman, S. Firestein, Q. Gong, P. D. Canoll, J. E. Goldman, P. Roussos, B. R. tenOever, B. O. Jonathan, S. Lomvardas, Non-cell-autonomous disruption of nuclear architecture as a potential cause of COVID-19-induced anosmia. Cell 185, 1052–1064 e1012 (2022).

9. B. D. Baxter, E. D. Larson, L. Merle, P. Feinstein, A. G. Polese, A. N. Bubak, C. S. Niemeyer, J. Hassell, Jr., D. Shepherd, V. R. Ramakrishnan, M. A. Nagel, D. Restrepo, Transcriptional profiling reveals potential involvement of microvillous TRPM5-expressing cells in viral infection of the olfactory epithelium. BMC Genomics 22, 224 (2021).

10. A. Hansen, T. E. Finger, Is TrpM5 a reliable marker for chemosensory cells? Multiple types of microvillous cells in the main olfactory epithelium of mice. BMC Neuroscience 9, 159–112 (2008).

11. W. Lin, E. A. Ezekwe, Jr., Z. Zhao, E. R. Liman, D. Restrepo, TRPM5-expressing microvillous cells in the main olfactory epithelium. BMC Neurosci 9, 114 (2008).

12. C. C. Hegg, C. Jia, W. S. Chick, D. Restrepo, A. Hansen, Microvillous cells expressing IP3 receptor type 3 in the olfactory epithelium of mice. Eur J Neurosci 32, 1632–1645 (2010).

13. S. Pfister, M. G. Dietrich, C. Sidler, J. M. Fritschy, I. Knuesel, R. Elsaesser, Characterization and turnover of CD73/IP(3)R3-positive microvillar cells in the adult mouse olfactory epithelium. Chem Senses 37, 859–868 (2012).

14. P. L. Weng, M. Vinjamuri, C. E. Ovitt, Ascl3 transcription factor marks a distinct progenitor lineage for non-neuronal support cells in the olfactory epithelium. Sci Rep 6, 38199 (2016).

15. C. Jia, S. Hayoz, C. R. Hutch, T. R. Iqbal, A. E. Pooley, C. C. Hegg, An IP3R3- and NPY-expressing microvillous cell mediates tissue homeostasis and regeneration in the mouse olfactory epithelium. PLoS One 8, e58668 (2013).

16. G. M. Goss, N. Chaudhari, J. M. Hare, R. Nwojo, B. Seidler, D. Saur, B. J. Goldstein, Differentiation potential of individual olfactory c-Kit+ progenitors determined via multicolor lineage tracing. Developmental Neurobiology 76, 241--251 (2016).

17. C. Jia, C. C. Hegg, NPY mediates ATP-induced neuroproliferation in adult mouse olfactory epithelium. Neurobiol Dis 38, 405–413 (2010).

18. K. Lemons, Z. Fu, I. Aoude, T. Ogura, J. Sun, J. Chang, K. Mbonu, I. Matsumoto, H. Arakawa, W. Lin, Lack of TRPM5-Expressing Microvillous Cells in Mouse Main Olfactory Epithelium Leads to Impaired Odor-Evoked Responses and Olfactory-Guided Behavior in a Challenging Chemical Environment. eNeuro 4, (2017).

19. T. Ogura, S. A. Szebenyi, K. Krosnowski, A. Sathyanesan, J. Jackson, W. Lin, Cholinergic microvillous cells in the mouse main olfactory epithelium and effect of acetylcholine on olfactory sensory neurons and supporting cells. J Neurophysiol 106, 1274–1287 (2011).

20. Z. Fu, T. Ogura, W. Luo, W. Lin, ATP and Odor Mixture Activate TRPM5-Expressing Microvillous Cells and Potentially Induce Acetylcholine Release to Enhance Supporting Cell Endocytosis in Mouse Main Olfactory Epithelium. Frontiers in Cellular Neuroscience 12, 10167–10116 (2018).

21. G. Krasteva, B. J. Canning, P. Hartmann, T. Z. Veres, T. Papadakis, C. Muhlfeld, K. Schliecker, Y. N. Tallini, A. Braun, H. Hackstein, N. Baal, E. Weihe, B. Schutz, M. Kotlikoff, I. Ibanez-Tallon, W. Kummer, Cholinergic chemosensory cells in the trachea regulate breathing. Proceedings of the National Academy of Sciences 108, 9478–9483 (2011).

22. S. Kaske, G. Krasteva, P. Konig, W. Kummer, T. Hofmann, T. Gudermann, V. Chubanov, TRPM5, a taste-signaling transient receptor potential ion-channel, is a ubiquitous signaling component in chemosensory cells. BMC Neurosci 8, 49 (2007).

23. C. Bezencon, A. Furholz, F. Raymond, R. Mansourian, S. Metairon, J. Le Coutre, S. Damak, Murine intestinal cells expressing Trpm5 are mostly brush cells and express markers of neuronal and inflammatory cells. J Comp Neurol 509, 514–525 (2008).

24. B. Schutz, I. Jurastow, S. Bader, C. Ringer, J. von Engelhardt, V. Chubanov, T. Gudermann, M. Diener, W. Kummer, G. Krasteva-Christ, E. Weihe, Chemical coding and chemosensory properties of cholinergic brush cells in the mouse gastrointestinal and biliary tract. Front Physiol 6, 87 (2015).

25. B. D. Gulbransen, T. R. Clapp, T. E. Finger, S. C. Kinnamon, Nasal Solitary Chemoreceptor Cell Responses to Bitter and Trigeminal Stimulants In Vitro. Journal of Neurophysiology 99, 2929–2937 (2008).

26. C. J. Saunders, M. Christensen, T. E. Finger, M. Tizzano, Cholinergic neurotransmission links solitary chemosensory cells to nasal inflammation. Proc Natl Acad Sci U S A 111, 6075–6080 (2014).

27. T. Yamaguchi, J. Yamashita, M. Ohmoto, I. Aoude, T. Ogura, W. Luo, A. A. Bachmanov, W. Lin, I. Matsumoto, J. Hirota, Skn-1a/Pou2f3 is required for the generation of Trpm5-expressing microvillous cells in the mouse main olfactory epithelium. BMC Neurosci 15, 13 (2014).

28. J. Yamashita, M. Ohmoto, T. Yamaguchi, I. Matsumoto, J. Hirota, Skn-1a/Pou2f3 functions as a master regulator to generate Trpm5-expressing chemosensory cells in mice. PLoS One 12, e0189340 (2017).

29. M. Ohmoto, T. Yamaguchi, J. Yamashita, A. A. Bachmanov, J. Hirota, I. Matsumoto, Pou2f3/Skn-1a is necessary for the generation or differentiation of solitary chemosensory cells in the anterior nasal cavity. Biosci Biotechnol Biochem 77, 2154–2156 (2013).

30. F. Gerbe, E. Sidot, D. J. Smyth, M. Ohmoto, I. Matsumoto, V. Dardalhon, P. Cesses, L. Garnier, M. Pouzolles, B. Brulin, M. Bruschi, Y. Harcus, V. S. Zimmermann, N. Taylor, R. M. Maizels, P. Jay, Intestinal epithelial tuft cells initiate type 2 mucosal immunity to helminth parasites. Nature 529, 226–230 (2016).

31. L. G. Bankova, D. F. Dwyer, E. Yoshimoto, S. Ualiyeva, J. W. McGinty, H. Raff, J. von Moltke, Y. Kanaoka, K. Frank Austen, N. A. Barrett, The cysteinyl leukotriene 3 receptor regulates expansion of IL-25-producing airway brush cells leading to type 2 inflammation. Science Immunology 3, eaat9453 (2018).

32. S. Ualiyeva, N. Hallen, Y. Kanaoka, C. Ledderose, I. Matsumoto, W. G. Junger, N. A. Barrett, L. G. Bankova, Airway brush cells generate cysteinyl leukotrienes through the ATP sensor P2Y2. Science immunology 5, (2020).

33. M. S. Nadjsombati, J. W. McGinty, M. R. Lyons-Cohen, J. B. Jaffe, L. DiPeso, C. Schneider, C. N. Miller, J. L. Pollack, G. A. N. Gowda, M. F. Fontana, D. J. Erle, M. S. Anderson, R. M. Locksley, D. Raftery, J. von Moltke, Detection of Succinate by Intestinal Tuft Cells Triggers a Type 2 Innate Immune Circuit. Immunity 49, 33–41.e37 (2018).

34. J. von Moltke, M. Ji, H.-E. Liang, R. M. Locksley, Tuft-cell-derived IL-25 regulates an intestinal ILC2–epithelial response circuit. Nature 529, 221–225 (2016).

35. A. L. Haber, M. Biton, N. Rogel, R. H. Herbst, K. Shekhar, C. Smillie, G. Burgin, T. M. Delorey, M. R. Howitt, Y. Katz, I. Tirosh, S. Beyaz, D. Dionne, M. Zhang, R. Raychowdhury, W. S. Garrett, O. Rozenblatt-Rosen, H. N. Shi, O. Yilmaz, R. J. Xavier, A. Regev, A Single-cell Survey of the Small Intestinal Epithelium. Nature 551, 333–339 (2017).

36. D. T. Montoro, A. L. Haber, M. Biton, V. Vinarsky, B. Lin, S. E. Birket, F. Yuan, S. Chen, H. M. Leung, J. Villoria, N. Rogel, G. Burgin, A. M. Tsankov, A. Waghray, M. Slyper, J. Waldman, L. Nguyen, D. Dionne, O. Rozenblatt-Rosen, P. R. Tata, H. Mou, M. Shivaraju, H. Bihler, M. Mense, G. J. Tearney, S. M. Rowe, J. F. Engelhardt, A. Regev, J. Rajagopal, A Revised Airway Epithelial Hierarchy Includes CFTR-expressing Ionocytes. Nature 560, 319–324 (2018).

37. M. R. Howitt, S. Lavoie, M. Michaud, A. M. Blum, S. V. Tran, J. V. Weinstock, C. A. Gallini, K. Redding, R. F. Margolskee, L. C. Osborne, D. Artis, W. S. Garrett, Tuft cells, taste-chemosensory cells, orchestrate parasite type 2 immunity in the gut. Science, 1–9 (2016).

38. T. E. Finger, B. Böttger, A. Hansen, K. T. Anderson, H. Alimohammadi, W. L. Silver, Solitary Chemoreceptor Cells in the Nasal Cavity Serve as Sentinels of Respiration. Proceedings of the National Academy of Sciences 100, 8981–8986 (2003).

39. F. Merigo, D. Benati, M. Tizzano, F. Osculati, A. Sbarbati, alpha-Gustducin immunoreactivity in the airways. Cell Tissue Res 319, 211–219 (2005).

40. F. Genovese, M. Tizzano, Microvillous Cells in the Olfactory Epithelium Express Elements of the Solitary Chemosensory Cell Transduction Signaling Cascade. PLoS one 13, e0202754–0202715 (2018).

41. S. Ualiyeva, E. Lemire, E. C. Aviles, C. Wong, A. A. Boyd, J. Lai, T. Liu, I. Matsumoto, N. A. Barrett, J. A. Boyce, A. L. Haber, L. G. Bankova, Tuft cell–produced cysteinyl leukotrienes and IL-25 synergistically initiate lung type 2 inflammation. Science Immunology 6, eabj0474 (2021).

42. J. Barr, M. E. Gentile, S. Lee, M. E. Kotas, M. Fernanda de Mello Costa, N. P. Holcomb, A. Jaquish, G. Palashikar, M. Soewignjo, M. McDaniel, I. Matsumoto, R. Margolskee, J. Von Moltke, N. A. Cohen, X. Sun, A. E. Vaughan, Injury-induced pulmonary tuft cells are heterogenous, arise independent of key Type 2 cytokines, and are dispensable for dysplastic repair. Elife 11, (2022).

43. C. K. Rane, S. R. Jackson, C. F. Pastore, G. Zhao, A. I. Weiner, N. N. Patel, D. R. Herbert, N. A. Cohen, A. E. Vaughan, Development of solitary chemosensory cells in the distal lung after severe influenza injury. American journal of physiology. Lung cellular and molecular physiology 316, L1141–L1149 (2019).

44. A. Ablimit, T. Aoki, T. Matsuzaki, T. Suzuki, H. Hagiwara, S. Takami, K. Takata, Immunolocalization of water channel aquaporins in the vomeronasal organ of the rat: expression of AQP4 in neuronal sensory cells. Chem Senses 33, 481–488 (2008).

45. P.-L. Weng, M. Vinjamuri, C. E. Ovitt, Ascl3 transcription factor marks a distinct progenitor lineage for non-neuronal support cells in the olfactory epithelium. Scientific Reports, 1–11 (2019).

46. S. Pfister, T. Weber, W. Härtig, C. Schwerdel, R. Elsaesser, I. Knuesel, J.-M. Fritschy, Novel role of cystic fibrosis transmembrane conductance regulator in maintaining adult mouse olfactory neuronal homeostasis. The Journal of comparative neurology 523, 406--430 (2015).

47. M. Tizzano, T. E. Finger, Chemosensors in the nose: guardians of the airways. Physiology (Bethesda) 28, 51–60 (2013).

48. R. J. Lee, N. A. Cohen, Sinonasal solitary chemosensory cells “taste” the upper respiratory environment to regulate innate immunity. Am J Rhinol Allergy 28, 366–373 (2014).

49. M. A. Kohanski, A. D. Workman, N. N. Patel, L. Y. Hung, J. P. Shtraks, B. Chen, M. Blasetti, L. Doghramji, D. W. Kennedy, N. D. Adappa, J. N. Palmer, D. R. Herbert, N. A. Cohen, Solitary chemosensory cells are a primary epithelial source of IL-25 in patients with chronic rhinosinusitis with nasal polyps. J Allergy Clin Immunol 142, 460–469 e467 (2018).

50. N. N. Patel, V. Triantafillou, I. W. Maina, A. D. Workman, C. C. L. Tong, E. C. Kuan, P. Papagiannopoulos, J. V. Bosso, N. D. Adappa, J. N. Palmer, M. A. Kohanski, D. R. Herbert, N. A. Cohen, Fungal extracts stimulate solitary chemosensory cell expansion in noninvasive fungal rhinosinusitis. Int Forum Allergy Rhinol 9, 730–737 (2019).

51. J. Ordovas-Montanes, D. F. Dwyer, S. K. Nyquist, K. M. Buchheit, M. Vukovic, C. Deb, M. H. Wadsworth, T. K. Hughes, S. W. Kazer, E. Yoshimoto, K. N. Cahill, N. Bhattacharyya, H. R. Katz, B. Berger, T. M. Laidlaw, J. A. Boyce, N. A. Barrett, A. K. Shalek, Allergic Inflammatory Memory in Human Respiratory Epithelial Progenitor Cells. Nature 560, 649–654 (2018).

52. D. T. Moran, J. C. Rowley, 3rd, B. W. Jafek, Electron microscopy of human olfactory epithelium reveals a new cell type: the microvillar cell. Brain Res 253, 39–46 (1982).

53. E. E. Morrison, R. M. Costanzo, Morphology of the human olfactory epithelium. J Comp Neurol 297, 1–13 (1990).

54. M. A. Durante, S. Kurtenbach, Z. B. Sargi, J. W. Harbour, R. Choi, S. Kurtenbach, G. M. Goss, H. Matsunami, B. J. Goldstein, Single-cell Analysis of Olfactory Neurogenesis and Differentiation in Adult Humans. Nature Neuroscience 23, 323–326 (2020).

55. F. Miragall, G. Kadmon, M. Husmann, M. Schachner, Expression of cell adhesion molecules in the olfactory system of the adult mouse: Presence of the embryonic form of N-CAM. Developmental Biology 129, 516--531 (1988).

56. R. A. Hall, Autonomic modulation of olfactory signaling. Science Signaling 4, pe1 (2011).

57. D. D. Gerendasy, J. G. Sutcliffe, RC3/neurogranin, a postsynaptic calpacitin for setting the response threshold to calcium influxes. Mol Neurobiol 15, 131–163 (1997).

58. F. Gerbe, B. Brulin, L. Makrini, C. Legraverend, P. Jay, DCAMKL-1 expression identifies Tuft cells rather than stem cells in the adult mouse intestinal epithelium. Gastroenterology 137, 2179–2180; author reply 2180-2171 (2009).

59. B. D. Gulbransen, T. E. Finger, Solitary chemoreceptor cell proliferation in adult nasal epithelium. J Neurocytol 34, 117–122 (2005).

60. M. Ohmoto, I. Matsumoto, A. Yasuoka, Y. Yoshihara, K. Abe, Genetic tracing of the gustatory and trigeminal neural pathways originating from T1R3-expressing taste receptor cells and solitary chemoreceptor cells. Mol Cell Neurosci 38, 505–517 (2008).

61. R. J. Lee, G. Xiong, J. M. Kofonow, B. Chen, A. Lysenko, P. Jiang, V. Abraham, L. Doghramji, N. D. Adappa, J. N. Palmer, D. W. Kennedy, G. K. Beauchamp, P. T. Doulias, H. Ischiropoulos, J. L. Kreindler, D. R. Reed, N. A. Cohen, T2R38 taste receptor polymorphisms underlie susceptibility to upper respiratory infection. J Clin Invest 122, 4145–4159 (2012).

62. J. D. Barros-Silva, D. E. Linn, I. Steiner, G. Guo, A. Ali, H. Pakula, G. Ashton, I. Peset, M. Brown, N. W. Clarke, R. T. Bronson, G. C. Yuan, S. H. Orkin, Z. Li, E. Baena, Single-Cell Analysis Identifies LY6D as a Marker Linking Castration-Resistant Prostate Luminal Cells to Prostate Progenitors and Cancer. Cell Rep 25, 3504–3518 e3506 (2018).

63. G. Upadhyay, Emerging Role of Lymphocyte Antigen-6 Family of Genes in Cancer and Immune Cells. Front Immunol 10, 819 (2019).

64. A. Mayama, K. Takagi, H. Suzuki, A. Sato, Y. Onodera, Y. Miki, M. Sakurai, T. Watanabe, K. Sakamoto, R. Yoshida, T. Ishida, H. Sasano, T. Suzuki, OLFM4, LY6D and S100A7 as potent markers for distant metastasis in estrogen receptor-positive breast carcinoma. Cancer Sci 109, 3350-3359 (2018).

65. A. M. Newman, C. B. Steen, C. L. Liu, A. J. Gentles, A. A. Chaudhuri, F. Scherer, M. S. Khodadoust, M. S. Esfahani, B. A. Luca, D. Steiner, M. Diehn, A. A. Alizadeh, Determining cell type abundance and expression from bulk tissues with digital cytometry. Nature Biotechnology 37, 773--782 (2019).

66. C. Schneider, C. E. O’Leary, R. M. Locksley, Regulation of immune responses by tuft cells. Nature Reviews Immunology, 1–10 (2019).

67. T. Kobayashi, K. Iijima, S. Radhakrishnan, V. Mehta, R. Vassallo, C. B. Lawrence, J. C. Cyong, L. R. Pease, K. Oguchi, H. Kita, Asthma-related environmental fungus, Alternaria, activates dendritic cells and produces potent Th2 adjuvant activity. Journal of immunology (Baltimore, Md. :1950) 182, 2502–2510 (2009).

68. H. Kouzaki, K. Iijima, T. Kobayashi, S. M. O’Grady, H. Kita, The danger signal, extracellular ATP, is a sensor for an airborne allergen and triggers IL-33 release and innate Th2-type responses. Journal of immunology (Baltimore, Md. : 1950) 186, 4375–4387 (2011).

69. T. A. Doherty, N. Khorram, J. E. Chang, H. K. Kim, P. Rosenthal, M. Croft, D. H. Broide, STAT6 regulates natural helper cell proliferation during lung inflammation initiated by Alternaria. American journal of physiology. Lung cellular and molecular physiology 303, L577–588 (2012).

70. T. A. Doherty, N. Khorram, K. Sugimoto, D. Sheppard, P. Rosenthal, J. Y. Cho, A. Pham, M. Miller, M. Croft, D. H. Broide, Alternaria induces STAT6-dependent acute airway eosinophilia and epithelial FIZZ1 expression that promotes airway fibrosis and epithelial thickness. Journal of immunology (Baltimore, Md. : 1950) 188, 2622–2629 (2012).

71. K. Ogawa, K. Asano, S. Yotsumoto, T. Yamane, M. Arita, Y. Hayashi, H. Harada, C. Makino-Okamura, H. Fukuyama, K. Kondo, T. Yamasoba, M. Tanaka, Frontline Science: Conversion of neutrophils into atypical Ly6G(+) SiglecF(+) immune cells with neurosupportive potential in olfactory neuroepithelium. J Leukoc Biol 109, 481–496 (2021).

72. T. A. Doherty, N. Khorram, S. Lund, A. K. Mehta, M. Croft, D. H. Broide, Lung type 2 innate lymphoid cells express cysteinyl leukotriene receptor 1, which regulates TH2 cytokine production. J Allergy Clin Immunol 132, 205–213 (2013).

73. C. S. Smillie, M. Biton, J. Ordovas-Montanes, K. M. Sullivan, G. Burgin, D. B. Graham, R. H. Herbst, N. Rogel, M. Slyper, J. Waldman, M. Sud, E. Andrews, G. Velonias, A. L. Haber, K. Jagadeesh, S. Vickovic, J. Yao, C. Stevens, D. Dionne, L. T. Nguyen, A.-C. Villani, M. Hofree, E. A. Creasey, H. Huang, O. Rozenblatt-Rosen, J. J. Garber, H. Khalili, A. N. Desch, M. J. Daly, A. N. Ananthakrishnan, A. K. Shalek, R. J. Xavier, A. Regev, Intra- and inter-cellular rewiring of the human colon during ulcerative colitis. Cell 178, 714–730.e722 (2019).

74. R. Vento-Tormo, M. Efremova, R. A. Botting, M. Y. Turco, M. Vento-Tormo, K. B. Meyer, J.-E. Park, E. Stephenson, K. Polanski, A. Goncalves, L. Gardner, S. Holmqvist, J. Henriksson, A. Zou, A. M. Sharkey, B. Millar, B. Innes, L. Wood, A. Wilbrey-Clark, R. P. Payne, M. A. Ivarsson, S. Lisgo, A. Filby, D. H. Rowitch, J. N. Bulmer, G. J. Wright, M. J. T. Stubbington, M. Haniffa, A. Moffett, S. A. Teichmann, Single-cell Reconstruction of the Early Maternal–fetal Interface in Humans. Nature 563, 347–353 (2018).

75. J. A. Ramilowski, T. Goldberg, J. Harshbarger, E. Kloppmann, E. Kloppman, M. Lizio, V. P. Satagopam, M. Itoh, H. Kawaji, P. Carninci, B. Rost, A. R. R. Forrest, A draft network of ligand-receptor-mediated multicellular signalling in human. Nature Communications 6, 7866 (2015).

76. B. J. Goldstein, G. M. Goss, K. E. Hatzistergos, E. B. Rangel, B. Seidler, D. Saur, J. M. Hare, Adult c-Kit(+) progenitor cells are necessary for maintenance and regeneration of olfactory neurons. J Comp Neurol 523, 15–31 (2015).

77. R. C. Krolewski, A. Packard, J. E. Schwob, Global expression profiling of globose basal cells and neurogenic progression within the olfactory epithelium. J Comp Neurol 521, 833–859 (2013).

78. J. E. Schwob, W. Jang, E. H. Holbrook, B. Lin, D. B. Herrick, J. N. Peterson, J. H. Coleman, Stem and progenitor cells of the mammalian olfactory epithelium: Taking poietic license. Journal of Comparative Neurology 525, 1034--1054 (2017).

79. T. T. Yu, J. C. McIntyre, S. C. Bose, D. Hardin, M. C. Owen, T. S. McClintock, Differentially expressed transcripts from phenotypically identified olfactory sensory neurons. J Comp Neurol 483, 251–262 (2005).

80. J. E. Schwob, S. L. Youngentob, R. C. Mezza, Reconstitution of the rat olfactory epithelium after methyl bromide-induced lesion. J Comp Neurol 359, 15–37 (1995).

81. M. Chen, R. R. Reed, A. P. Lane, Acute inflammation regulates neuroregeneration through the NF-kappaB pathway in olfactory epithelium. Proceedings of the National Academy of Sciences 114, 8089–8094 (2017).

82. M. Chen, R. R. Reed, A. P. Lane, Chronic Inflammation Directs an Olfactory Stem Cell Functional Switch from Neuroregeneration to Immune Defense. Cell Stem Cell 25, 501--513.e505 (2019).

83. T. T. Solbu, T. Holen, Aquaporin pathways and mucin secretion of Bowman’s glands might protect the olfactory mucosa. Chemical Senses 37, 35--46 (2012).

84. R. B. Fletcher, D. Das, L. Gadye, K. N. Street, A. Baudhuin, A. Wagner, M. B. Cole, Q. Flores, Y. G. Choi, N. Yosef, E. Purdom, S. Dudoit, D. Risso, J. Ngai, Deconstructing Olfactory Stem Cell Trajectories at Single-Cell Resolution. Cell Stem Cell 20, 817–830.e818 (2017).

85. T. Oguma, K. Asano, A. a. Ishizaka, Role of prostaglandin D2 and its receptors in the pathophysiology of asthma. Allergology International 57, 307–312 (2008).

86. A. S. Evans, N. J. Lennemann, C. B. Coyne, BPIFB3 interacts with ARFGAP1 and TMED9 to regulate non-canonical autophagy and RNA virus infection. Journal of Cell Science 134, jcs251835 (2021).

87. A. S. Evans, N. J. Lennemann, C. B. Coyne, BPIFB3 Regulates Endoplasmic Reticulum Morphology To Facilitate Flavivirus Replication. Journal of Virology 94, e00029–00020 (2020).

88. S. Morosky, N. J. Lennemann, C. B. Coyne, BPIFB6 Regulates Secretory Pathway Trafficking and Enterovirus Replication. J Virol 90, 5098–5107 (2016).

89. D. M. Irwin, J. M. Biegel, C. B. Stewart, Evolution of the mammalian lysozyme gene family. BMC Evol Biol 11, 166 (2011).

90. S. Mukherjee, C. L. Partch, R. E. Lehotzky, C. V. Whitham, H. Chu, C. L. Bevins, K. H. Gardner, L. V. Hooper, Regulation of C-type lectin antimicrobial activity by a flexible N-terminal prosegment. J Biol Chem 284, 4881–4888 (2009).

91. W. M. Weston, E. E. LeClair, W. Trzyna, K. M. McHugh, P. Nugent, C. M. Lafferty, L. Ma, R. S. Tuan, R. M. Greene, Differential display identification of plunc, a novel gene expressed in embryonic palate, nasal epithelium, and adult lung. J Biol Chem 274, 13698–13703 (1999).

92. M. Musa, K. Wilson, L. Sun, A. Mulay, L. Bingle, H. M. Marriott, E. E. LeClair, C. D. Bingle, Differential localisation of BPIFA1 (SPLUNC1) and BPIFB1 (LPLUNC1) in the nasal and oral cavities of mice. Cell Tissue Res 350, 455–464 (2012).

93. L. Bingle, S. S. Cross, A. S. High, W. A. Wallace, D. A. Devine, S. Havard, M. A. Campos, C. D. Bingle, SPLUNC1 (PLUNC) is expressed in glandular tissues of the respiratory tract and in lung tumours with a glandular phenotype. J Pathol 205, 491–497 (2005).

94. J. Pevsner, P. M. Hwang, P. B. Sklar, J. C. Venable, S. H. Snyder, Odorant-binding protein and its mRNA are localized to lateral nasal gland implying a carrier function. Proceedings of the National Academy of Sciences 85, 2383–2387 (1988).

95. M. Tegoni, P. Pelosi, F. Vincent, S. Spinelli, V. Campanacci, S. Grolli, R. Ramoni, C. Cambillau, Mammalian Odorant Binding Proteins. Biochimica et Biophysica Acta 1482, 229–240 (2000).

96. T. E. Sutherland, N. Logan, D. Rckerl, A. A. Humbles, S. M. Allan, V. Papayannopoulos, B. Stockinger, R. M. Maizels, J. E. Allen, Chitinase-like proteins promote IL-17-mediated neutrophilia in a tradeoff between nematode killing and host damage. Nature Immunology 15, 1116–1125 (2014).

97. A. Saraswathula, M. M. Liu, H. Kulaga, A. P. Lane, Chronic interleukin-13 expression in mouse olfactory mucosa results in regional aneuronal epithelium. Int Forum Allergy Rhinol 13, 230–241 (2023).

98. V. A. Epstein, P. J. Bryce, D. B. Conley, R. C. Kern, A. M. Robinson, Intranasal Aspergillus fumigatus exposure induces eosinophilic inflammation and olfactory sensory neuron cell death in mice. Otolaryngol Head Neck Surg 138, 334–339 (2008).

99. N. Schnittke, D. B. Herrick, B. Lin, J. Peterson, J. H. Coleman, A. I. Packard, W. Jang, J. E. Schwob, Transcription factor p63 controls the reserve status but not the stemness of horizontal basal cells in the olfactory epithelium. Proceedings of the National Academy of Sciences 112, E5068--E5077 (2015).

100. R. B. Fletcher, M. S. Prasol, J. Estrada, A. Baudhuin, K. Vranizan, Y. G. Choi, J. Ngai, p63 regulates olfactory stem cell self-renewal and differentiation. Neuron 72, 748--759 (2011).

101. A. Packard, N. Schnittke, R. A. Romano, S. Sinha, J. E. Schwob, Np63 Regulates Stem Cell Dynamics in the Mammalian Olfactory Epithelium. Journal of Neuroscience 31, 8748--8759 (2011).

102. D. B. Herrick, B. Lin, J. Peterson, N. Schnittke, J. E. Schwob, Notch1 maintains dormancy of olfactory horizontal basal cells, a reserve neural stem cell. Proceedings of the National Academy of Sciences 114, E5589--E5598 (2017).

103. F. A. Wolf, F. K. Hamey, M. Plass, J. Solana, J. S. Dahlin, B. Gttgens, N. Rajewsky, L. Simon, F. J. Theis, PAGA: graph abstraction reconciles clustering with trajectory inference through a topology preserving map of single cells. Genome Biology 20, 59 (2019).

104. X. Wang, N. R. Hallen, M. Lee, S. Samuchiwal, Q. Ye, K. M. Buchheit, A. Z. Maxfield, R. E. Roditi, R. W. Bergmark, N. Bhattacharyya, T. Ryan, D. Gakpo, S. Raychaudhuri, D. Dwyer, T. M. Laidlaw, J. A. Boyce, M. Gutierrez-Arcelus, N. A. Barrett, Type 2 Inflammation Drives an Airway Basal Stem Cell Program Through Insulin Receptor Substrate Signaling. J Allergy Clin Immunol, (2023).

105. A. M. Dittrich, M. Krokowski, H. A. Meyer, D. Quarcoo, A. Avagyan, B. Ahrens, S. M. Kube, M. Witzenrath, C. Loddenkemper, J. B. Cowland, E. Hamelmann, Lipocalin2 protects against airway inflammation and hyperresponsiveness in a murine model of allergic airway disease. Clin Exp Allergy 40, 1689–1700 (2010).

106. N. Zimmermann, M. P. Doepker, D. P. Witte, K. F. Stringer, P. C. Fulkerson, S. M. Pope, E. B. Brandt, A. Mishra, N. E. King, N. M. Nikolaidis, M. Wills-Karp, F. D. Finkelman, M. E. Rothenberg, Expression and regulation of small proline-rich protein 2 in allergic inflammation. Am J Respir Cell Mol Biol 32, 428–435 (2005).

107. J. J. Peschon, P. J. Morrissey, K. H. Grabstein, F. J. Ramsdell, E. Maraskovsky, B. C. Gliniak, L. S. Park, S. F. Ziegler, D. E. Williams, C. B. Ware, J. D. Meyer, B. L. Davison, Early lymphocyte expansion is severely impaired in interleukin 7 receptor-deficient mice. J Exp Med 180, 1955–1960 (1994).

108. M. H. Kaplan, U. Schindler, S. T. Smiley, M. J. Grusby, Stat6 is required for mediating responses to IL-4 and for development of Th2 cells. Immunity 4, 313–319 (1996).

109. M. S. Kowalczyk, I. Tirosh, D. Heckl, T. N. Rao, A. Dixit, B. J. Haas, R. K. Schneider, A. J. Wagers, B. L. Ebert, A. Regev, Single-cell RNA-seq reveals changes in cell cycle and differentiation programs upon aging of hematopoietic stem cells. Genome Research 25, 1860–1872 (2015).

110. C. E. O’Leary, C. Schneider, R. M. Locksley, Tuft Cells-Systemically Dispersed Sensory Epithelia Integrating Immune and Neural Circuitry. Annu Rev Immunol, (2018).

111. C. D. Brown, I. Kilty, M. Yeadon, S. Jenkinson, Regulation of 15-lipoxygenase isozymes and mucin secretion by cytokines in cultured normal human bronchial epithelial cells. Inflamm Res 50, 321–326 (2001).

112. P. G. Woodruff, H. A. Boushey, G. M. Dolganov, C. S. Barker, Y. H. Yang, S. Donnelly, A. Ellwanger, S. S. Sidhu, T. P. Dao-Pick, C. Pantoja, D. J. Erle, K. R. Yamamoto, J. V. Fahy, Genome-wide profiling identifies epithelial cell genes associated with asthma and with treatment response to corticosteroids. Proc Natl Acad Sci U S A 104, 15858–15863 (2007).

113. K. Lemons, Z. Fu, T. Ogura, W. Lin, TRPM5-expressing Microvillous Cells Regulate Region-specific Cell Proliferation and Apoptosis During Chemical Exposure. Neuroscience 434, 171--190 (2020).

114. K. F. Austen, A. Maekawa, Y. Kanaoka, J. A. Boyce, The leukotriene E4 puzzle: finding the missing pieces and revealing the pathobiologic implications. J Allergy Clin Immunol 124, 406–414; quiz 415-406 (2009).

115. C. B. Westphalen, S. Asfaha, Y. Hayakawa, Y. Takemoto, D. J. Lukin, A. H. Nuber, A. Brandtner, W. Setlik, H. Remotti, A. Muley, X. Chen, R. May, C. W. Houchen, J. G. Fox, M. D. Gershon, M. Quante, T. C. Wang, Long-lived intestinal tuft cells serve as colon cancer-initiating cells. J Clin Invest 124, 1283–1295 (2014).

116. K. Deckmann, K. Filipski, G. Krasteva-Christ, M. Fronius, M. Althaus, A. Rafiq, T. Papadakis, L. Renno, I. Jurastow, L. Wessels, M. Wolff, B. Schutz, E. Weihe, V. Chubanov, T. Gudermann, J. Klein, T. Bschleipfer, W. Kummer, Bitter triggers acetylcholine release from polymodal urethral chemosensory cells and bladder reflexes. Proc Natl Acad Sci U S A 111, 8287–8292 (2014).

117. S. Ualiyeva, E. Yoshimoto, N. A. Barrett, L. G. Bankova, Isolation and Quantitative Evaluation of Brush Cells from Mouse Tracheas. J Vis Exp 59496, (2019).

118. B. Lin, J. H. Coleman, J. N. Peterson, M. J. Zunitch, W. Jang, D. B. Herrick, J. E. Schwob, Injury Induces Endogenous Reprogramming and Dedifferentiation of Neuronal Progenitors to Multipotency. Cell Stem Cell 21, 761–774 e765 (2017).

119. T. Alejandre-Garcia, J. G. Pena-Del Castillo, A. Hernandez-Cruz, GABAA receptor: a unique modulator of excitability, Ca(2+) signaling, and catecholamine release of rat chromaffin cells. Pflugers Arch 470, 67-77 (2018).

120. J. Peterson, B. Lin, C. M. Barrios-Camacho, D. B. Herrick, E. H. Holbrook, W. Jang, J. H. Coleman, J. E. Schwob, Activating a Reserve Neural Stem Cell Population In Vitro Enables Engraftment and Multipotency after Transplantation. Stem Cell Reports 12, 680–695 (2019).

121. N. Schaefer, X. Li, M. A. Seibold, N. N. Jarjour, L. C. Denlinger, M. Castro, A. M. Coverstone, W. G. Teague, J. Boomer, E. R. Bleecker, D. A. Meyers, W. C. Moore, G. A. Hawkins, J. Fahy, B. R. Phillips, D. T. Mauger, A. Dakhama, S. Gellatly, N. Pavelka, R. Berman, Y. P. Di, S. E. Wenzel, H. W. Chu, The effect of BPIFA1/SPLUNC1 genetic variation on its expression and function in asthmatic airway epithelium. JCI Insight 4, e127237 (2019).

122. T. Wu, J. Huang, P. J. Moore, M. S. Little, W. G. Walton, R. C. Fellner, N. E. Alexis, Y. P. Di, M. R. Redinbo, S. L. Tilley, R. Tarran, Identification of BPIFA1/SPLUNC1 as an epithelium-derived smooth muscle relaxing factor. Nature Communications 8, 14118 (2017).

123. Y. Enomoto, K. Orihara, T. Takamasu, A. Matsuda, Y. Gon, H. Saito, C. Ra, Y. Okayama, Tissue remodeling induced by hypersecreted epidermal growth factor and amphiregulin in the airway after an acute asthma attack. Journal of Allergy and Clinical Immunology 124, 913--920.e917 (2009).

124. Y. Zhu, X. Yan, C. Zhai, L. Yang, M. Li, Association between risk of asthma and gene polymorphisms in CHI3L1 and CHIA: a systematic meta-analysis. BMC Pulmonary Medicine 17, 193 (2017).

125. K. Wu, K. Kamimoto, Y. Zhang, K. Yang, S. P. Keeler, B. J. Gerovac, E. V. Agapov, S. P. Austin, J. Yantis, K. A. Gissy, D. E. Byers, J. Alexander-Brett, C. M. Hoffmann, M. Wallace, M. E. Hughes, E. C. Crouch, S. A. Morris, M. J. Holtzman, Basal-epithelial stem cells cross an alarmin checkpoint for post-viral lung disease. Journal of Clinical Investigation 131, (2021).

126. M. E. Kotas, C. M. Moore, J. G. Gurrola, 2nd, S. D. Pletcher, A. N. Goldberg, R. Alvarez, S. Yamato, P. E. Bratcher, C. A. Shaughnessy, P. L. Zeitlin, I. H. Zhang, Y. Li, M. T. Montgomery, K. Lee, E. K. Cope, R. M. Locksley, M. A. Seibold, E. D. Gordon, IL-13-programmed airway tuft cells produce PGE2, which promotes CFTR-dependent mucociliary function. JCI Insight 7, (2022).

127. K. M. Buchheit, A. Sohail, J. Hacker, R. Maurer, D. Gakpo, J. C. Bensko, F. Taliaferro, J. Ordovas-Montanes, T. M. Laidlaw, Rapid and sustained effect of dupilumab on clinical and mechanistic outcomes in aspirin-exacerbated respiratory disease. J Allergy Clin Immunol 150, 415–424 (2022).

128. G. Finak, A. McDavid, M. Yajima, J. Deng, V. Gersuk, A. K. Shalek, C. K. Slichter, H. W. Miller, M. J. McElrath, M. Prlic, P. S. Linsley, R. Gottardo, MAST: a flexible statistical framework for assessing transcriptional changes and characterizing heterogeneity in single-cell RNA sequencing data. Genome Biology 16, 1–13 (2015).

129. L. Scrucca, M. Fop, T. B. Murphy, A. E. Raftery, mclust 5: Clustering, Classification and Density Estimation Using Gaussian Finite Mixture Models. The R Journal 8, 289–317 (2016).

130. F. A. Wolf, P. Angerer, F. J. Theis, SCANPY: large-scale single-cell gene expression data analysis. Genome Biology 19, 1–5 (2018).

131. V. A. Traag, L. Waltman, N. J. van Eck, From Louvain to Leiden: guaranteeing well-connected communities. Scientific Reports 9, 5233 (2019).

132. L. Albergante, E. Mirkes, J. Bac, H. Chen, A. Martin, L. Faure, E. Barillot, L. Pinello, A. Gorban, A. Zinovyev, Robust and Scalable Learning of Complex Intrinsic Dataset Geometry via ElPiGraph. Entropy 22, 296 (2020).

133. S. N. Wood, Fast stable restricted maximum likelihood and marginal likelihood estimation of semiparametric generalized linear models. Journal of the Royal Statistical Society B 73, 3--36 (2011).

134. J. X. Binder, S. Pletscher-Frankild, K. Tsafou, C. Stolte, S. I. O’Donoghue, R. Schneider, L. J. Jensen, COMPARTMENTS: unification and visualization of protein subcellular localization evidence. Database: the journal of biological databases and curation 2014, bau012 (2014).

135. B. Li, C. N. Dewey, RSEM: accurate transcript quantification from RNA-Seq data with or without a reference genome. BMC Bioinformatics 12, 323 (2011).

136. P.-C. Bürkner, brms: An R Package for Bayesian Multilevel Models Using Stan. Journal of Statistical Software 80, 1–28 (2017).

